# Outdoor Air Pollution, Perivascular Space Morphology, and Cognition in Preadolescence

**DOI:** 10.1101/2025.09.26.678867

**Authors:** Jessica Morrel, Kirthana Sukumaran, Carinna Torgerson, Michael Rosario, Haoyu Lan, Joel Schwartz, Jiu-Chiuan Chen, Jeiran Choupan, Megan M. Herting

## Abstract

**Background:** Ambient air pollution exposure is associated with structural brain differences and poorer cognition in children; however, mechanisms of toxicity remain unclear. Perivascular spaces (PVS), key for brain waste clearance, may play a role in the neurotoxicity of air pollution. This study explored associations between air pollution exposure, PVS morphology, and cognition in preadolescents.

**Methods:** We analyzed cross-sectional Adolescent Brain Cognitive Development^SM^ (ABCD) Study^®^ data from 6,949 9-10-year-old participants. Annual average exposures to PM_2.5_, O_3_, NO_2_, and 15 PM_2.5_ components were estimated using spatiotemporal models mapped to residential addresses. PVS count and volume were derived from T1w and T2w MRI, and cognition was estimated using NIH Toolbox scores. Linear mixed-effects models examined independent associations between air pollution, PVS, and cognition; weighted quantile sum regression assessed co-exposure effects of PM_2.5_ mixtures.

**Findings:** Linear models revealed that exposures to Zn, NH_4_ ^+^, and Br were positively associated with PVS count in several regions. Higher PVS count in five key regions was associated with poorer cognitive performance across several NIH Toolbox domains. Higher Ca, Zn, and NH _4_ ^+^ exposures were associated with poorer cognition (P_FDR_ < 0.01). Higher frontal lobe PVS count mediated the association between Zn exposure and poorer total cognition (P < 0.01). Co-exposure models revealed that PM_2.5_ mixtures were associated with higher temporal and cingulate PVS count, and poorer working memory and crystallized intelligence (P < 0.01).

**Interpretation:** Outdoor air pollution was associated with higher PVS count and reduced cognition, suggesting that brain clearance may be a novel mechanism linking pollution to neurodevelopmental harm in preadolescents.

**Funding:** This work was supported by the National Institutes of Health (NIH) National Institute of Environmental Health Sciences (NIEHS) (Grant Nos. R01ES032295 and R01ES031074 [to MMH]; T32ES013678 [to JM]; P30ES07048 [to JM and MAR]; 3P30ES000002-55S [to MAR]), National Institute of Mental Health (NIMH) (Grant RF1MH123223 [to JC]), National Institute of Neurological Disorders and Stroke (Grant R01NS128486 [to JC]), and EPA grants (Grant Nos. 83587201 and 83544101 [to JS]).

## 1. Introduction

Outdoor air pollution is a ubiquitous neurotoxicant associated with a host of structural and functional brain differences^1^. Mounting evidence supports a link between prenatal and childhood air pollution exposure, particularly to PM_2.5_ and its components, and poor cognition, including worse inhibitory control^2,3^, processing speed^4,5^, attention^5,6^, working memory^3,7,8^, and verbal intelligence^9^. While animal studies implicate microglial activation, inflammation, white matter injury, and alterations in synaptic plasticity^10^, specific mechanisms by which air pollution impacts the developing brain and cognition remain unclear. Here, we investigate brain perivascular spaces (PVS) as one potential target of air pollution induced neurotoxicity.

PVS are cerebrospinal fluid (CSF)-filled spaces surrounding cerebral vasculature (**Figure 1A**) and a key component of the brain’s glymphatic system, which is responsible for facilitating nutrient delivery, waste clearance, and implicated in neuroinflammation and immune cell trafficking^11–13^. PVS can be visualized using magnetic resonance imaging (MRI) (**Figure 1B**) and quantified by the number visible on an MRI (count), by their volume, or other morphological and functional properties^14^. Studies demonstrate that MRI-visible PVS are present from infancy and increase in count and volume across the lifespan^15–17^. Considering the high metabolic demands of neurodevelopment^18^, efficient nutrient transport and waste clearance are necessary, making the PVS of particular interest during childhood and adolescence.

**Figure 1.**
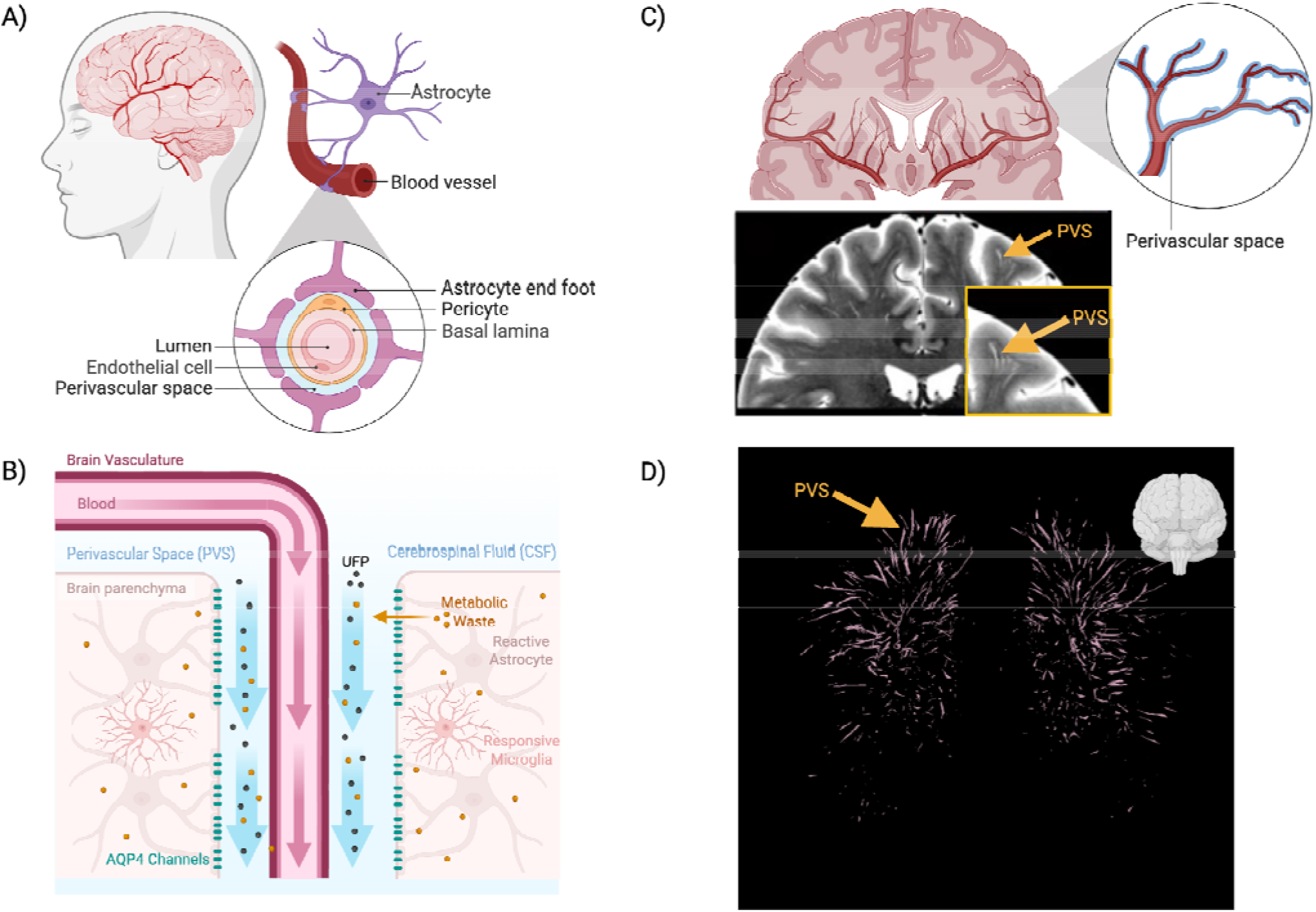
Conceptual schematic and 3-dimensional, in-vivo perivascular space (PVS) segmentations from a representative ABCD Study participant. **A)** Schematic depicting PVS anatomy, **B)** Schematic depicting PVS waste clearance flow and hypothesized route of pollution-induced toxicity, **C**) Conceptual and in-vivo rendering of PVS on a T2-weighted magnetic resonance imaging (MRI) scan, **D)** 3-dimensional, in-vivo PVS segmentations generated using the Weakly Supervised Perivascular Spaces Segmentation (WPSS) pipeline from the frontal view. 3-dimensional rendering created using the Quantitative Imaging Toolbox^46^. Abbreviations: Perivascular Space (PVS); Ultrafine Particles (UFP); Aquaporin-4 (AQP4). Image created using Biorender.com.

Evidence suggests a link between enlarged PVS, reduced glymphatic clearance, and poorer cognition in older adults^19,20^. Mechanistically, there are several hypotheses. PVS play an important role in neurovascular coupling^21^, which has been positively associated with cognition^22,23^. Furthermore, PVS are a site of immune cell accumulation^24^ and a key actor in neuroinflammatory processes^12^. Therefore, in the presence of an immunological challenge, such as air pollution exposure, it is plausible that PVS may be a site of neurotoxicity^25^. Findings from one study using light and electron microscopy in dogs, children, and teens found that exposure to O_3_ and PM_2.5_ was associated with perivascular and neurovascular unit dysfunction in the prefrontal white matter; authors note this may be a mechanism of Alzheimer’s Disease (AD) pathogenesis^26^. While clinical studies have examined cognitive symptoms in hospitalized children with enlarged PVS^27,28^ and two studies have examined air pollution and PVS in adults^29,30^, none, to our knowledge, has tested the relationship between air pollution exposure, PVS morphology, and cognitive functioning in a large group of healthy children.

The current study fills this gap by exploring novel relationships between air pollution exposure, PVS count (number of MRI-visible spaces) and volume fraction (VF), and cognition in 9-10-year-olds. Using linear mixed-effects (LME) models and weighted quantile sum (WQS) regression, we examined relationships between exposure and co-exposure to three criteria pollutants (PM_2.5_, NO_2_, O_3_) and 15 PM_2.5_ components, PVS morphology, and cognition. Considering that enlarged PVS are rarely observed in healthy children^31^, and that PM_2.5_ and metal components are associated with oxidative stress and inflammation^32^, we hypothesized that exposure to PM_2.5_ and its metal components would be associated with higher MRI-visible PVS count. Next, we expected that higher PVS count would be negatively associated with memory and processing speed, considering the evidence linking PVS burden to these two cognitive outcomes^19^. Finally, we expected that PVS morphology would mediate the relationship between PM_2.5_ and cognition.

## 2. Methods

### 2.1 Study Design

This study used cross-sectional imaging, air pollution, and cognitive data collected during the baseline visit of the ongoing Adolescent Brain Cognitive Development^SM^ (ABCD) Study^□^ (NIMH Data Archive annual 3.0 [imaging] and 5.0 [all other data] releases; DOI: 10.15154/8873-zj6, DOI: 10.15154/1520591). The ABCD study enrolled 11,880 9-10-year-old children (mean age = 9.49 years; 48% female) from 21 sites across the United States between October 2016 and October 2018^33–35^. Centralized Institutional Review Board (IRB) approval was obtained from the University of California San Diego; each site obtained approval from their local IRB. Written consent was provided by the child’s parent or legal guardian, and written assent was obtained from each child. Eligibility requirements for enrollment included 9.0-10.99 years at the baseline visit and English fluency. Exclusion criteria included severe sensory, neurological, medical, or intellectual limitations, and inability to complete the MRI scan. Participants were included in the current study if they had residentially mapped air pollution data at baseline and passed all neuroimaging and perivascular space (PVS) segmentation quality control criteria (**Section 2.3**). 6,949 participants were included in the current study (see **Figure S1** for flowchart of participant exclusion).

### 2.2 Air Pollution Exposure

Procedures for estimating annual average concentrations of ambient outdoor air pollution exposure have been previously described^36,37^. Daily exposures to PM_2.5_ (µg/m^3^), NO_2_ (ppb), and the 8-hour daily maximums for O_3_ (ppb) were calculated at 1-km^2^ resolution using hybrid spatiotemporal modeling that incorporates satellite-based aerosol optical depth models, land-use regression, meteorological data, and chemical transport model outputs^38,39^. Monthly concentrations for 15 PM_2.5_ components—bromine (Br), calcium (Ca), copper (Cu), elemental carbon (EC), iron (Fe), potassium (K), ammonium (NH_4_ ^+^), nickel (Ni), nitrate (NO_3_ ^-^), organic carbon (OC), lead (Pb), silicon (Si), sulfate (SO_4_ ^2-^), vanadium (V), zinc (Zn)—were estimated at a 50-meter resolution using similar hybrid modeling methods^40^. Exposure estimates for criteria pollutants and PM_2.5_ components were averaged across the 2016 calendar year, corresponding to the baseline enrollment visit, and were assigned to each child’s primary residential address at baseline (**Table S1**).

### 2.3 Neuroimaging Data

Raw T1w and T2w images meeting quality control standards were downloaded from the ABCD 3.0 release. Due to the sensitivity of PVS pipelines to motion and artifacts, only participants passing both ABCD raw image and Freesurfer quality control (QC) were included in the current study (**Figure S1**). For PVS segmentation, raw T1w and T2w images were registered via the Human Connectome Project (HCP) minimal preprocessing pipeline^41^, where data were upsampled to 0.7 mm^3^ resolution to enhance signal contrast. Preprocessed T1w and T2w images were filtered using adaptive non-local mean filtering^42^ and the enhanced PVS contrast (EPC) was obtained by dividing the filtered T1w image/T2w image. EPC images were generated to enhance PVS visibility^43^. Finally, all EPC images underwent manual visual QC to remove participants with extreme noise or incidental findings. Next, the Weakly Supervised Perivascular Spaces Segmentation was applied as previously described^44^. See **Supplement Section A.1** for additional details. 3-dimensional, in-vivo segmentations of PVS for a representative participant are visualized in **Figure 1C, 1D**. The white matter was then parcellated using the Desikan-Killiany atlas^45^ and pooled into six key regions of interest–frontal, temporal, parietal, occipital, cingulate, and centrum semiovale (CSO). PVS volume and count were calculated for these six regions. For PVS volumes, volume fractions (VF) were calculated by dividing the volume of PVS in each region by that region’s total volume, to normalize PVS volume between regions by their size.

### 2.4 Cognitive Data

The NIH Toolbox^□^ cognitive battery was collected at the baseline study visit, including tasks that assess language skills and receptive vocabulary (Picture Vocabulary Task), expressive language and reading decoding skills (Oral Reading Recognition Task), rapid visual processing speed (Pattern Comparison Processing Speed Test), working memory (List Sorting Working Memory Test), episodic memory (Picture Sequence Memory Test), inhibitory control and attention (Flanker Task), and cognitive flexibility (Dimensional Change Card Sort Task)^47^. From these tasks, three summary scores are derived: Total Cognition Composite, Fluid Composite (includes Dimensional Change Card Sort, Flanker, Picture

Sequence Memory, List Sorting Working Memory, and Pattern Comparison Tasks), and Crystallized Composite (includes Picture Vocabulary and Oral Reading Recognition Tasks). We utilized age-corrected scores^48^.

### 2.5 Demographics & Covariates

Covariates were selected based on previous literature and the construction of directed acyclic graphs (**Figure S2**)^49^. Sociodemographic variables such as parent-reported race/ethnicity (*White, Black, Hispanic, Asian*, or *Other*) and average household income in US Dollars (*<50K, 50-100K, >100K, Don’t Know/Refuse to Answer*), were included due to known differences in air pollution exposure across socioeconomic and racial/ethnic groups^50,51^. We also included demographic factors variables which are known to be associated with PVS morphology, such as age at MRI, sex (*female* or *male*), and body mass index (z-scored; BMIz)^15,17,52^. Precision imaging variables included regional white matter volume in analyses with PVS count as the main outcome and a combination variable of MRI model and head coil (*32-* or *64-channel*), to account for PVS differences attributable to variability in cranial size and scanner differences. Finally, ABCD site was included as a random effect to account for the hierarchical nature of the ABCD Study.

### 2.6 Statistical Analyses

All data preparation and analyses were conducted using *R* Version 4.3.2^53^. Three sets of single-pollutant linear mixed-effects (LME) models were run to test independent associations between 1) air pollution and PVS morphology, 2) PVS count and cognition, and 3) air pollution and cognition. First, 36 models (analysis 1) per region (i.e., frontal, temporal, parietal, occipital, CSO, and cingulate) were performed to test associations between each pollutant (i.e., three criteria pollutants, 15 PM_2.5_ components) and PVS count or VF. Analyses using PVS count as the outcome assess whether exposure to air pollution is associated with a greater number of detectable PVS, reflecting potential increases in the frequency of small, MRI-visible PVS, while analyses using VF evaluate whether air pollution is linked to increased cumulative PVS burden within a region, which may reflect dilation or enlargement of individual PVS, independent of their number. Next, due to widespread associations seen between air pollution and PVS count, but not VF, we ran 60 post-hoc LME models (analysis 2) testing associations between PVS count in the six regions and the 10 cognitive outcomes. For analysis 3, we tested independent associations between 18 air pollutants and 10 NIH Toolbox outcomes and performed mediation analysis (see **Supplemental Methods A.3.4** for details) to determine whether PVS morphology mediates the relationship between air pollution exposure and cognition. False discovery rate (FDR) correction was implemented and only results passing FDR correction for significance at P_FDR_ < 0.01 were considered for mediation analysis. Cohen’s effect size was calculated, as well.

Finally, because pollutant exposure never occurs in isolation, we examined the relationships between co-exposure to the 15 PM_2.5_ components and PVS count, PVS VF, and cognition using weighted quantile sum (WQS) regression. WQS uses a generalized linear regression framework to model the exposure outcome relationship^54^ and creates a composite mixture index in which each component is assigned a data-driven weight, reflecting its importance in the association between the exposure of mixture index and outcome (see **Supplemental Methods A.2**). Separate WQS models were constructed for each outcome of interest, including PVS count and VF across six key regions, and 10 cognitive outcomes. Directionality of the associations were set a priori based on the observed trends from the LME results (i.e., positive for PVS, negative for cognition). All results presented were adjusted for the previously mentioned minimally sufficient set of confounders (**Section 2.5**). Component weights above 0.06, calculated based on the commonly used threshold, the inverse of the number of components (1/15)^54^, were considered important contributors to the WQS index. See **Supplemental Methods A.3** for model equations.

## 3. Results

Sociodemographic characteristics of the final study sample are included in **Table 1** and **Table S2**. Compared to the entire cohort, the current ABCD sample is slightly older and more likely to be female, white, and scanned on a Siemens MRI scanner (Ps < 0.05 - 0.001; **Table S2**). Participants were also exposed to marginally more O_3_ and less NO _4_^+^ (Ps < 0.001), NO_3_ ^-^, and SO _4_^2-^ (Ps < 0.05) than the whole ABCD cohort (see **Table S3** for descriptives**; Figures S3-S4** for correlograms). PVS count ranged from 124.2 in the occipital lobe to 1,029.7 in the CSO, on average. VFs remained relatively consistent across regions, ranging from 0.02 in the occipital lobe to 0.006 in the frontal lobe and CSO (see **Table S4; Figures S5-S6** for correlograms).

**Table 1.** Sample characteristics for the study sample. Total number (N) of participants and (percentage of total sample) for each demographic category are presented for the final study sample. Mean and standard deviation (SD) are presented for numeric variables. Abbreviations: Adolescent Brain Cognitive Development (ABCD) Study; General Educational Development (GED); General Electric (GE); Body Mass Index (BMI).

|  | N (% of Sample) |
| --- | --- |
| Total N | 6,949 (58.7%) |
| Sex (F) | 3,414 (49.1%) |
| *Race & Ethnicity* |  |
| Asian | 161 (2.3%) |
| Hispanic | 1,456 (21.0%) |
| Black | 920 (13.2%) |
| White | 3,720 (53.5%) |
| Other | 691 (9.9%) |
| *Household Income* |  |
| < \$50,000 | 1,779 (25.6%) |
| ≥\$50,000 to <\$100,000 | 1,846 (26.6%) |
| ≥ \$100,000 | 2,746 (39.5%) |
| Don't Know / Refuse | 578 (8.3%) |
| *Caregiver Education* |  |
| < High school diploma | 318 (4.6%) |
| High school diploma / GED | 632 (9.1%) |
| Some college, associate's degree | 1,737 (25.0%) |
| Bachelor's degree | 1,776 (25.6%) |
| Graduate degree | 2,481 (35.7%) |
| *MRI Scanner Manufacturer* |  |
| Siemens | 4,705 (67.7%) |
| GE Medical Systems | 1,429 (20.6%) |
| Philips Medical Systems | 815 (11.7%) |
| Mean (SD) |  |
| Age at baseline (months) | 119.2 (7.5) |
| BMI Z-Score | 0.4 (1.2) |

### 3.1 Single-Pollutant Models

#### 3.1.1 Analysis 1. Air Pollution and PVS Morphology

Several significant positive associations were seen between ambient air pollution exposure and PVS count and VF (P_FDR_ < 0.01) (**Figure 2; Tables 2, S5**). The strongest associations included higher Br exposure with increased PVS count in the cingulate, as well as higher Zn exposure with increased PVS counts in the frontal lobe and CSO. Higher NH_4_ was also associated with larger temporal PVS VF.

**Table 2.**
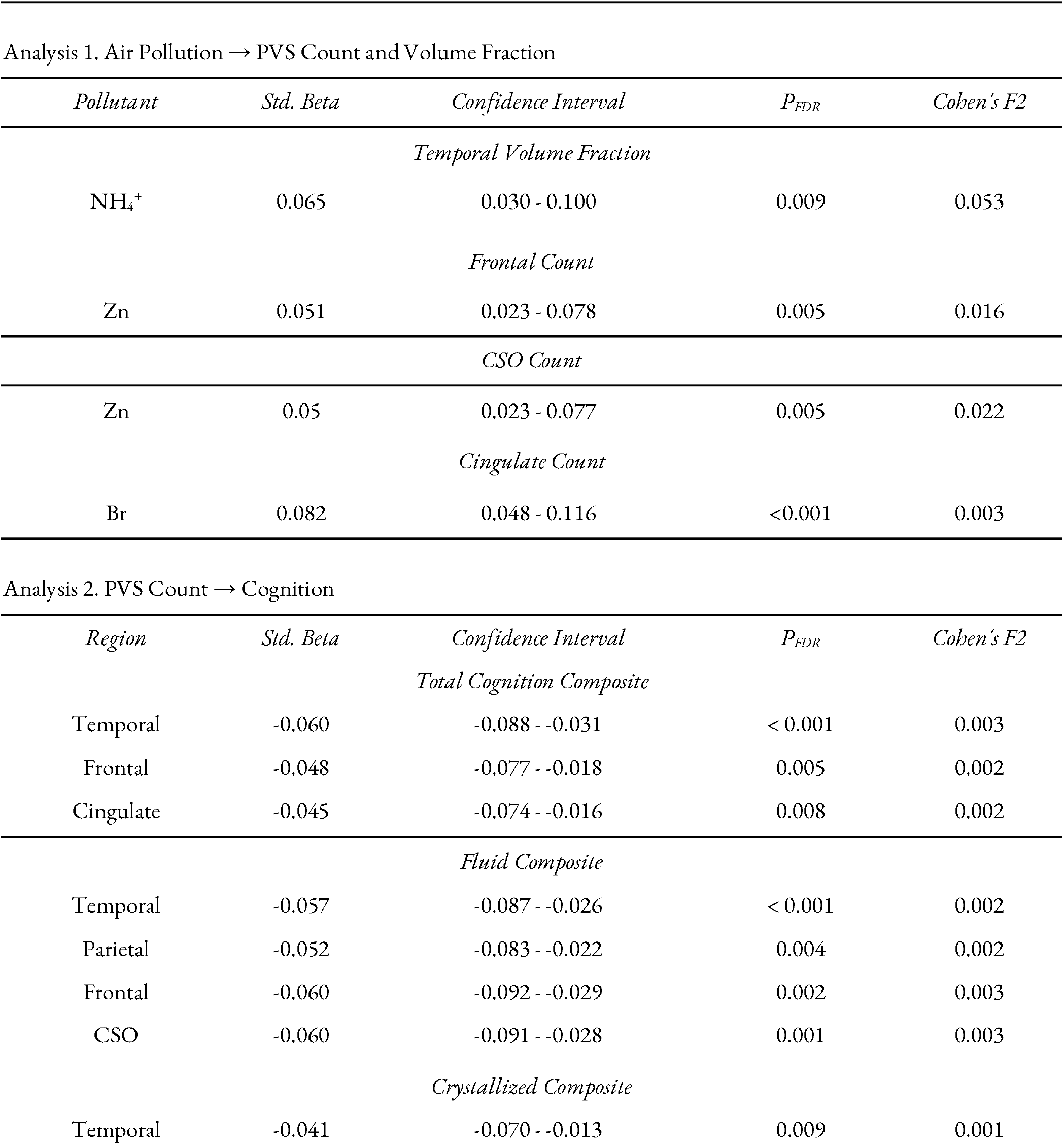
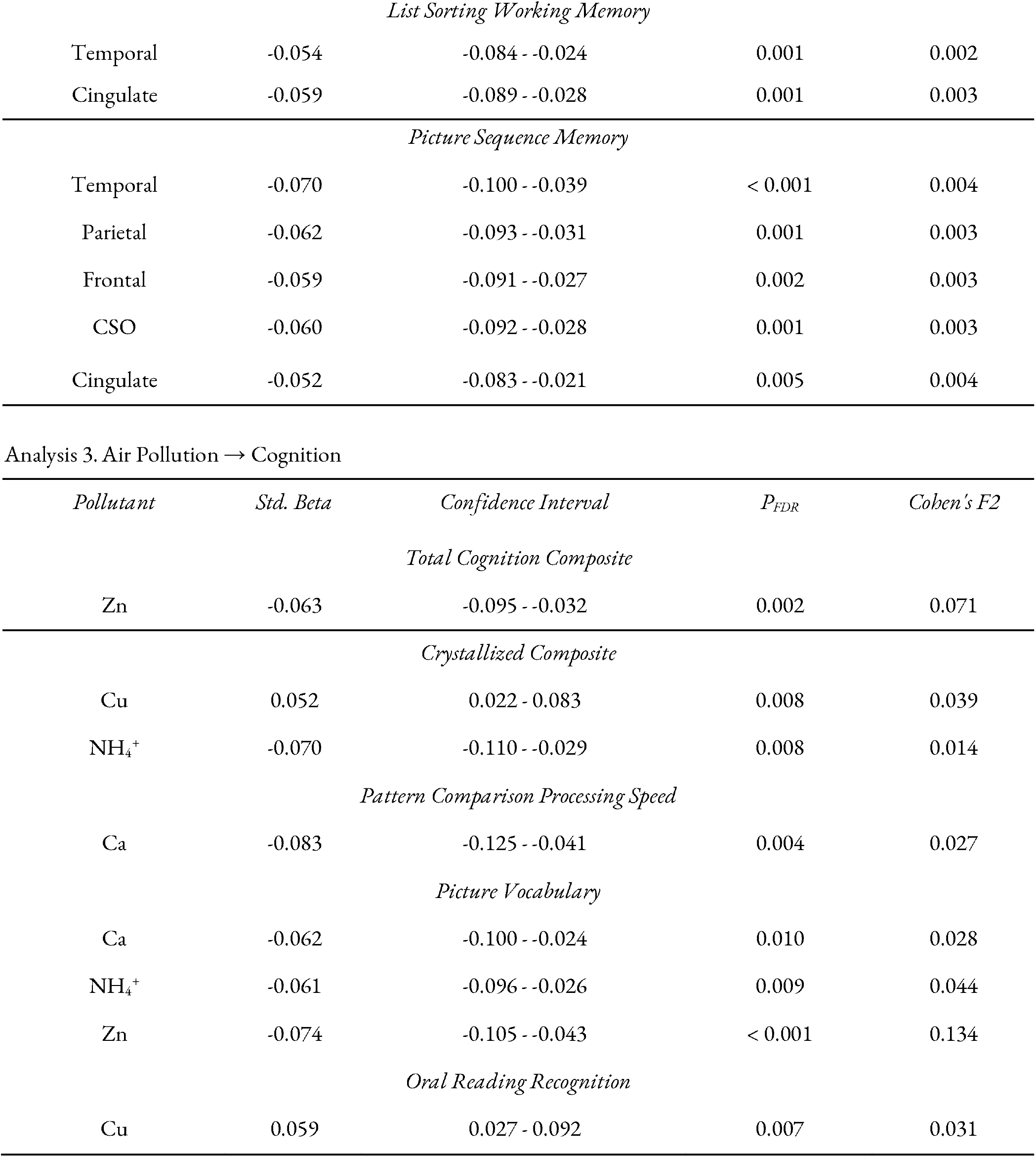
Results from single-pollutant linear mixed-effects models (p_FDR_ < 0.01) examining relationships between air pollution exposure, PVS count and volume fraction, and cognition. Abbreviations: Standardized (Std.); Perivascular space (PVS); Fine Particulate Matter (PM_2.5_), Nitrogen Dioxide (NO_2_), Ozone (O_3_), Bromine (Br), Calcium (Ca), Copper (Cu), Elemental Carbon (EC), Iron (Fe), Potassium (K), Ammonium (NH_4_^+^), Nitrate (NO_3_^-^), Nickel (Ni), Organic Carbon (OC), Lead (Pb), Sulfate (SO_4_^2-^), Silicon (Si), Vanadium (V), Zinc (Zn); False Discovery Rate (FDR); Centrum Semiovale (CSO).

| *Pollutant* | *Std. Beta* | *Confidence Interval* | *P<sub>FDR</sub>* | *Cohen's F2* |
| --- | --- | --- | --- | --- |
| *Temporal Volume Fraction* |  |  |  |  |
| NH<sub>4</sub><sup>+</sup> | 0.065 | 0.030 - 0.100 | 0.009 | 0.053 |
| *Frontal Count* |  |  |  |  |
| Zn | 0.051 | 0.023 - 0.078 | 0.005 | 0.016 |
| *CSO Count* |  |  |  |  |
| Zn | 0.05 | 0.023 - 0.077 | 0.005 | 0.022 |
| *Cingulate Count* |  |  |  |  |
| Br | 0.082 | 0.048 - 0.116 | <0.001 | 0.003 |

Analysis 2. PVS Count → Cognition
| *Region* | *Std. Beta* | *Confidence Interval* | *P<sub>FDR</sub>* | *Cohen's F2* |
| --- | --- | --- | --- | --- |
| *Total Cognition Composite* |  |  |  |  |
| Temporal | -0.060 | -0.088 - -0.031 | < 0.001 | 0.003 |
| Frontal | -0.048 | -0.077 - -0.018 | 0.005 | 0.002 |
| Cingulate | -0.045 | -0.074 - -0.016 | 0.008 | 0.002 |
| *Fluid Composite* |  |  |  |  |
| Temporal | -0.057 | -0.087 - -0.026 | < 0.001 | 0.002 |
| Parietal | -0.052 | -0.083 - -0.022 | 0.004 | 0.002 |
| Frontal | -0.060 | -0.092 - -0.029 | 0.002 | 0.003 |
| CSO | -0.060 | -0.091 - -0.028 | 0.001 | 0.003 |
| *Crystallized Composite* |  |  |  |  |
| Temporal | -0.041 | -0.070 - -0.013 | 0.009 | 0.001 |
| *List Sorting Working Memory* |  |  |  |  |
| Temporal | -0.054 | -0.084 - -0.024 | 0.001 | 0.002 |
| Cingulate | -0.059 | -0.089 - -0.028 | 0.001 | 0.003 |
| *Picture Sequence Memory* |  |  |  |  |
| Temporal | -0.070 | -0.100 - -0.039 | < 0.001 | 0.004 |
| Parietal | -0.062 | -0.093 - -0.031 | 0.001 | 0.003 |
| Frontal | -0.059 | -0.091 - -0.027 | 0.002 | 0.003 |
| CSO | -0.060 | -0.092 - -0.028 | 0.001 | 0.003 |
| Cingulate | -0.052 | -0.083 - -0.021 | 0.005 | 0.004 |

Analysis 3. Air Pollution → Cognition
| *Pollutant* | *Std. Beta* | *Confidence Interval* | *P<sub>FDR</sub>* | *Cohen's F2* |
| --- | --- | --- | --- | --- |
| *Total Cognition Composite* |  |  |  |  |
| Zn | -0.063 | -0.095 - -0.032 | 0.002 | 0.071 |
| *Crystallized Composite* |  |  |  |  |
| Cu | 0.052 | 0.022 - 0.083 | 0.008 | 0.039 |
| NH<sub>4</sub><sup>+</sup> | -0.070 | -0.110 - -0.029 | 0.008 | 0.014 |
| *Pattern Comparison Processing Speed* |  |  |  |  |
| Ca | -0.083 | -0.125 - -0.041 | 0.004 | 0.027 |
| *Picture Vocabulary* |  |  |  |  |
| Ca | -0.062 | -0.100 - -0.024 | 0.010 | 0.028 |
| NH<sub>4</sub><sup>+</sup> | -0.061 | -0.096 - -0.026 | 0.009 | 0.044 |
| Zn | -0.074 | -0.105 - -0.043 | < 0.001 | 0.134 |
| *Oral Reading Recognition* |  |  |  |  |
| Cu | 0.059 | 0.027 - 0.092 | 0.007 | 0.031 |

**Figure 2.**
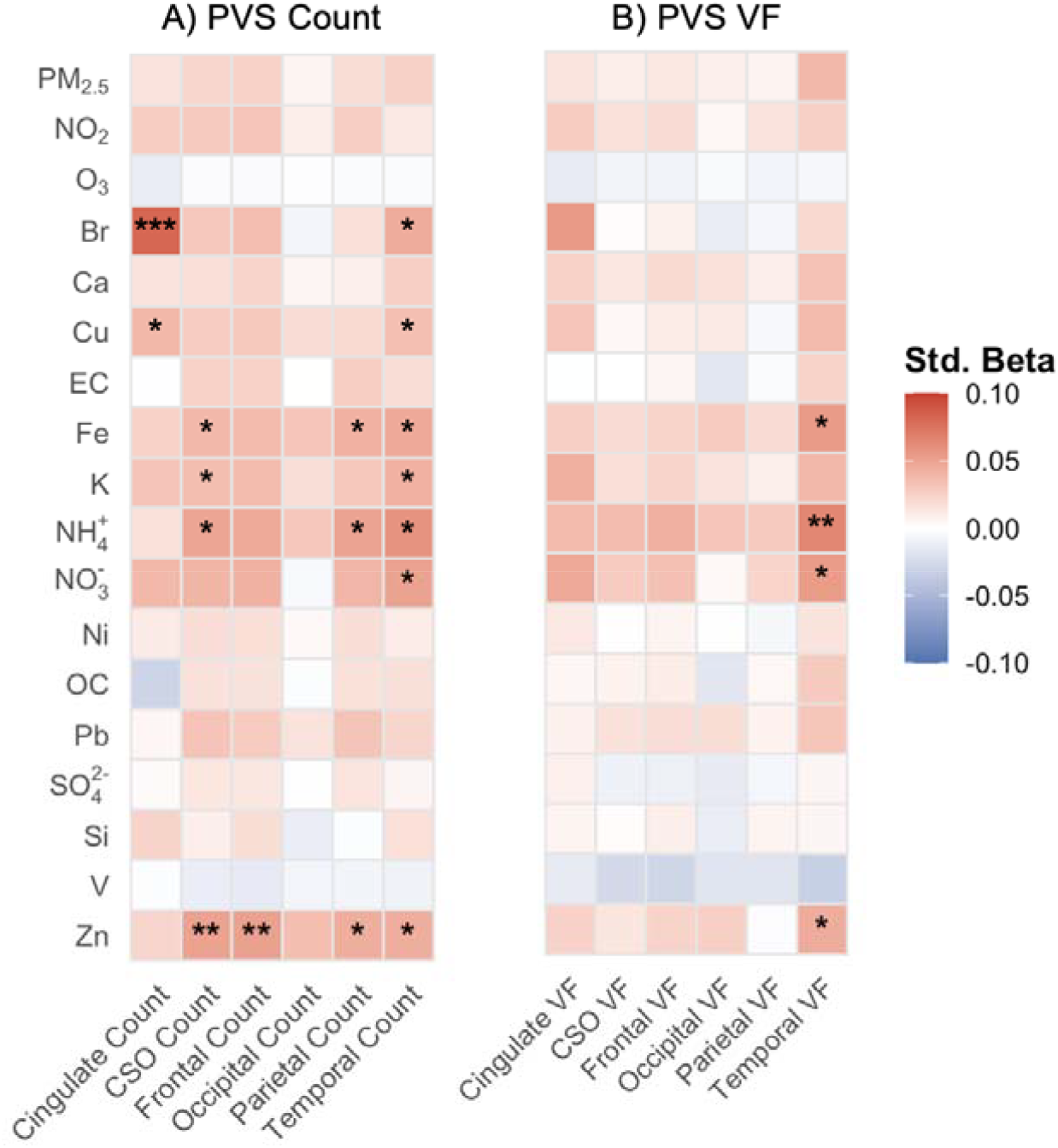
Associations between Air Pollution Exposure and PVS A) Count, and B) Volume Fraction (VF). Heatmap reflects standardized betas; stars reflect significance threshold where * = P_FDR_ < 0.05, ** = P_FDR_ < 0.01, and *** = P_FDR_ < 0.001. Abbreviations: Perivascular Spaces (PVS); Fine Particulate Matter (PM_2.5_), Nitrogen Dioxide (NO_2_), Ozone (O_3_), Bromine (Br), Calcium (Ca), Copper (Cu), Elemental Carbon (EC), Iron (Fe), Potassium (K), Ammonium (NH_4_ ^+^), Nitrate (NO_3_ ^-^), Nickel (Ni), Organic Carbon (OC), Lead (Pb), Sulfate (SO_4_ ^2-^), Silicon (Si), Vanadium (V), Zinc (Zn); Centrum Semiovale (CSO); Volume Fraction (VF); Standardized (Std).

#### 3.1.2 Analysis 2. PVS Count and Cognition

We identified significant negative associations between PVS count and several cognitive domains (P_FDR_ < 0.01) (**Figure 3; Tables 2, S6**). Higher PVS count in the temporal, frontal and cingulate was associated with poorer cognitive performance overall (i.e., total composite). Higher PVS count in the temporal, frontal, parietal, and CSO were also associated with lower fluid intelligence, whereas higher PVS count in only the temporal lobe was associated with poorer crystalized intelligence. Higher PVS counts in all regions, except the occipital lobe, were related to learning and memory performance. Contrary to hypotheses, significant associations were not observed for pattern comparison processing speed at P_FDR_ < 0.01, though associations were seen at P_FDR_ < 0.05.

**Figure 3.**
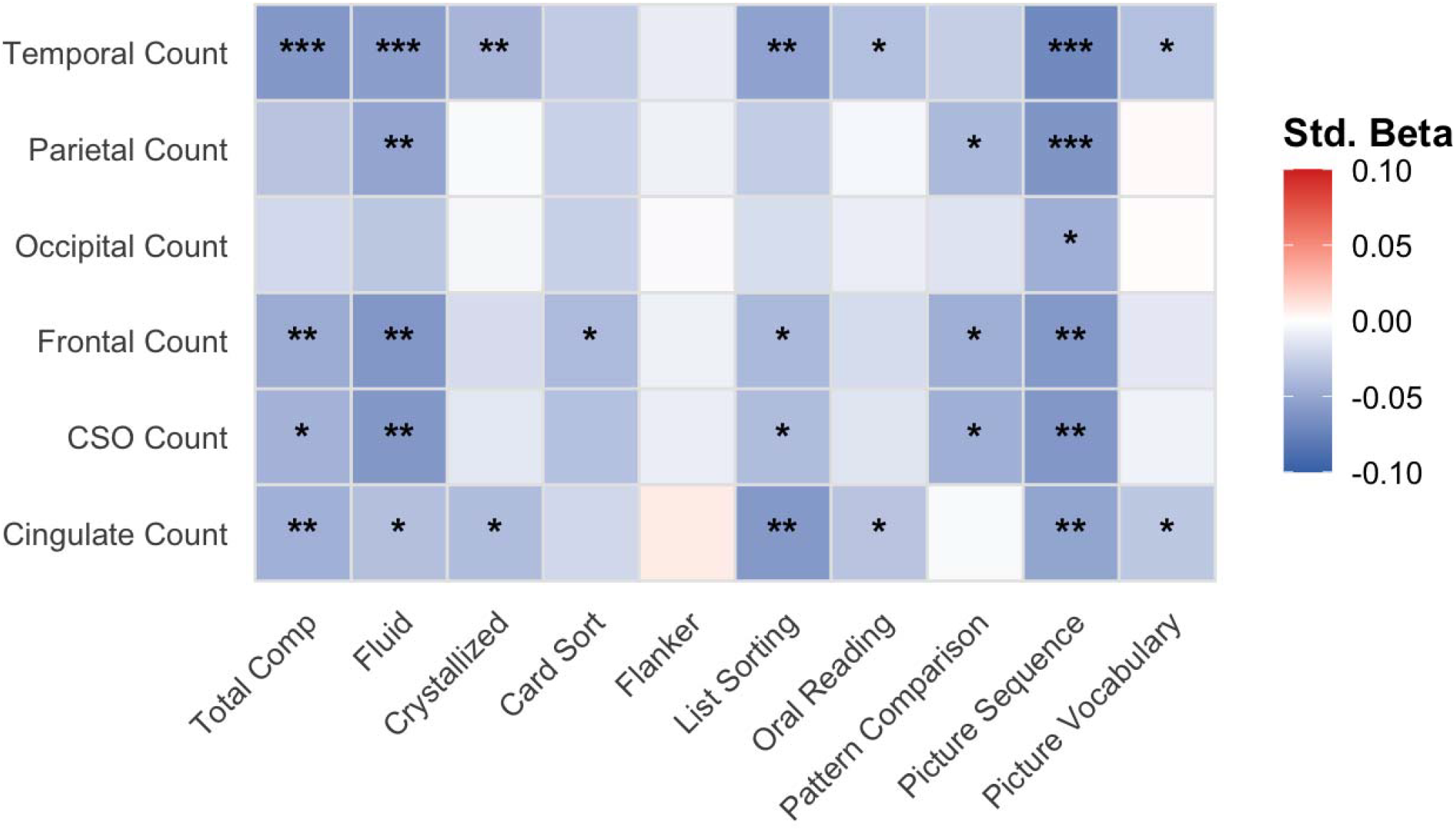
Associations between PVS Count and Cognition. Heatmap reflects standardized betas; stars r flect significance threshold where * = P_FDR_ < 0.05, ** = P_FDR_ < 0.01, and *** = P_FDR_ < 0.001. Abbreviations: Centrum Semiovale (CSO); Standardized (Std).

#### 3.1.3 Analysis 3. Mediation Analyses

After observing results suggesting a potential mediating pathway, we conducted a second set of single-pollutant regressions exploring the associations between air pollution exposure and cognition (P_FDR_ < 0.01) (**Figure 4; Table 2, S7**). Results from these models were mixed. While, contrary to hypotheses, Cu exposure was associated with higher oral reading recognition and crystallized composite scores, all other significant results demonstrated expected negative associations between air pollution exposure and cognitive measures. In particular, Ca exposure was associated with lower picture vocabulary and pattern comparison processing speed scores; NH_4_ ^+^ was negatively associated with picture vocabulary and crystallized composite scores; Zn was associated with lower picture vocabulary and total cognition composite scores.

**Figure 4.**
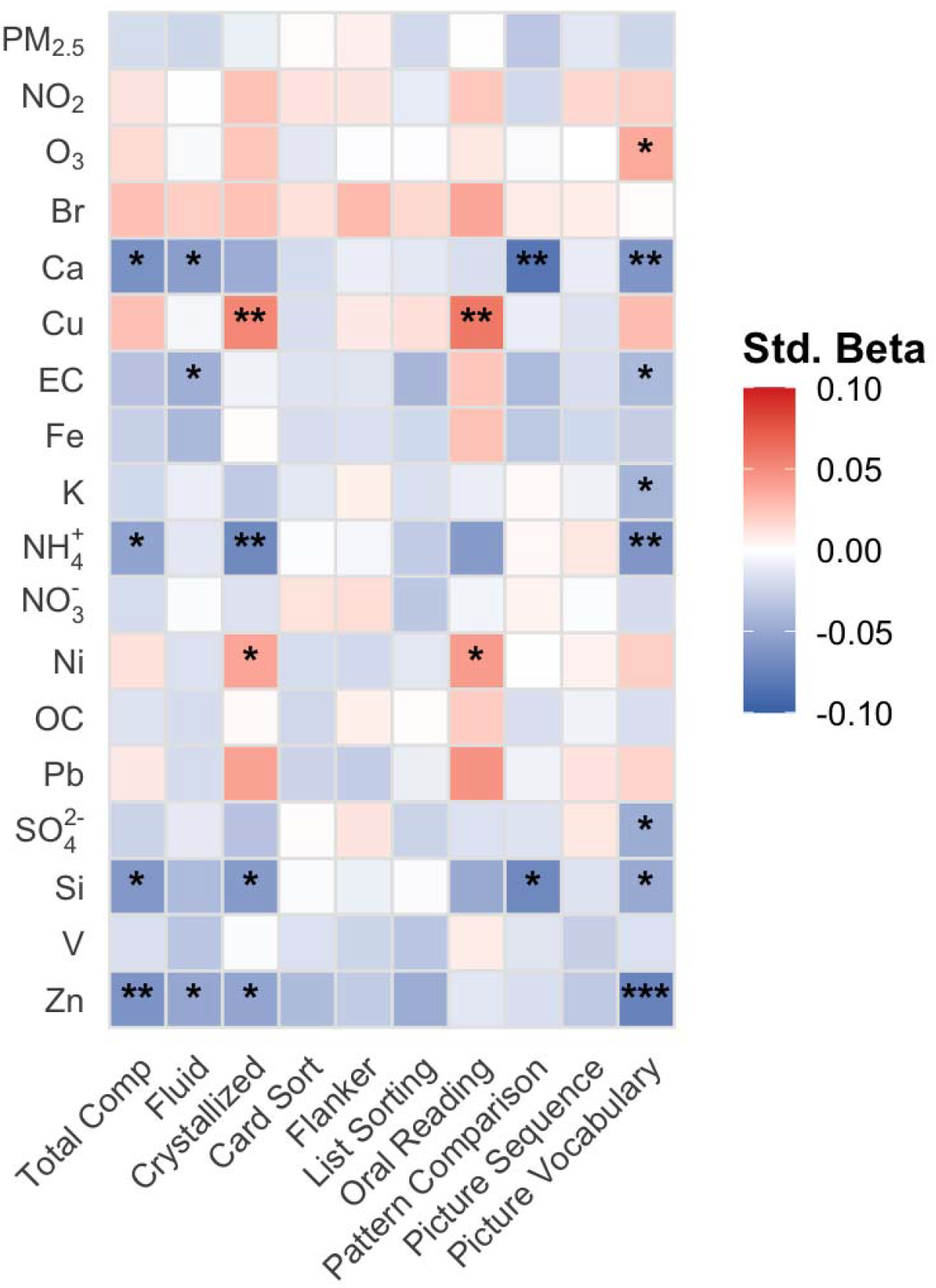
Associations between Air Pollution Exposure and Cognition. Heatmap reflects standardized betas; stars reflect significance threshold where * = P_FDR_ < 0.05, ** = P_FDR_ < 0.01, and *** = P_FDR_ < 0.001. Abbreviations: Fine Particulate Matter (PM_2.5_), Nitrogen Dioxide (NO_2_), Ozone (O_3_), Bromine (Br), Calcium (Ca), Copper (Cu), Elemental Carbon (EC), Iron (Fe), Potassium (K), Ammonium (NH_4_ ^+^), Nitrate (NO_3_ ^-^), Nickel (Ni), Organic Carbon (OC), Lead (Pb), Sulfate (SO_4_ ^2-^), Silicon (Si), Vanadium (V), Zinc (Zn); Standardized (Std).

In post-hoc testing of significant results (P_FDR_ < 0.01), we tested one potential mediation and detected a significant effect using the Baron and Kenny method^55^ and LME for mediation analysis, where PVS count in the frontal lobe mediated the negative association between Zn and total cognition composite (**Figure 5**).

**Figure 5.**
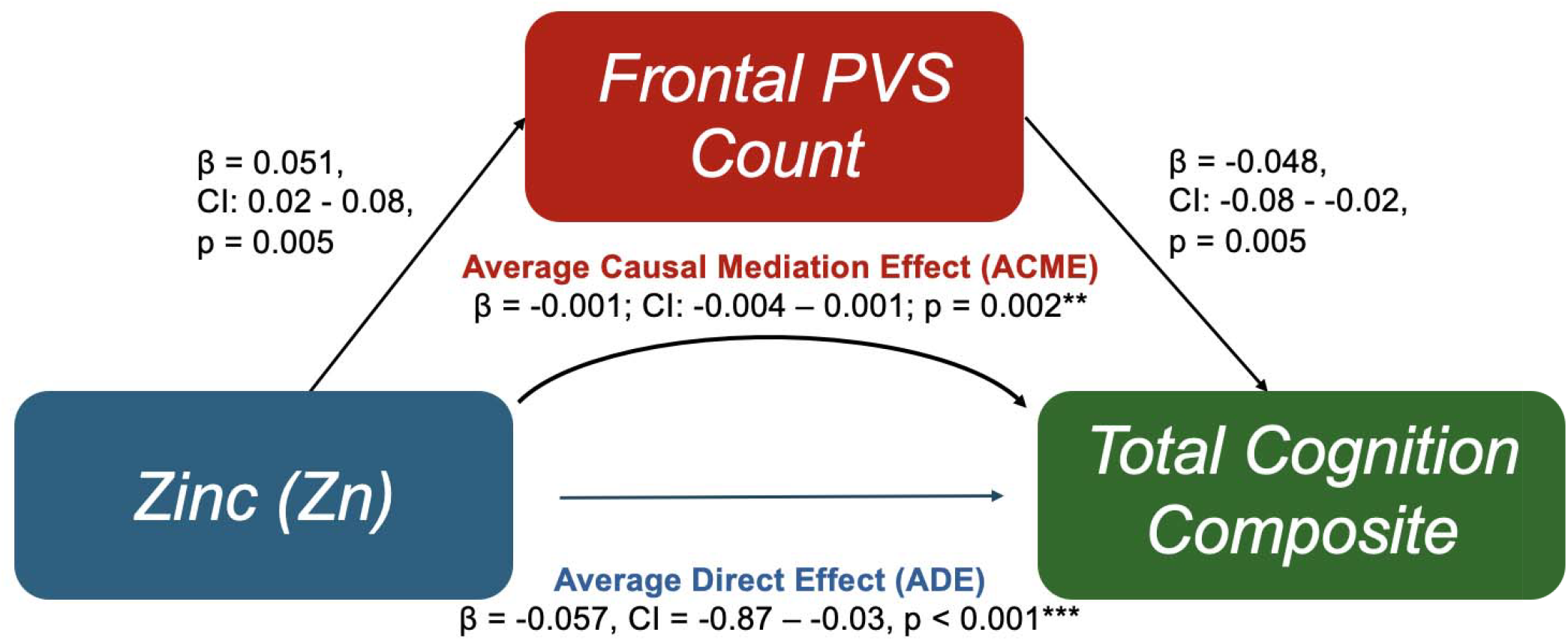
Mediation results depicting relationships between zinc (Zn), frontal PVS count, and total cogn tion composite. Abbreviations: Confidence Interval (CI); Perivascular Space (PVS).

### 3.2 Multi-Pollutant Models

#### 3.2.1 Analysis 1. Air Pollution and PVS morphology

WQS models examining associations between co-exposure to the 15 PM_2.5_ components and PVS count and VF yielded two significant results. Specifically, co-exposure was associated with higher temporal (P < 0.01) and cingulate (P < 0.001) PVS count (**Table 3, Figure S7, Table S8)**. Applying a weight cutoff of 0.06, NO_3_ ^-^, Br, Si, Cu, and NH_4_ ^+^ drove associations with temporal PVS count; the same components, along with V, drove associations with cingulate PVS count.

**Table 3.**
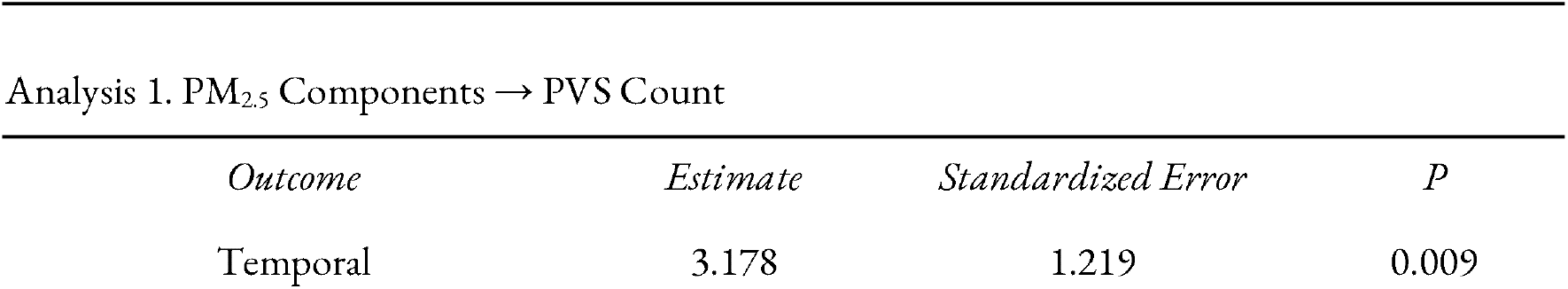
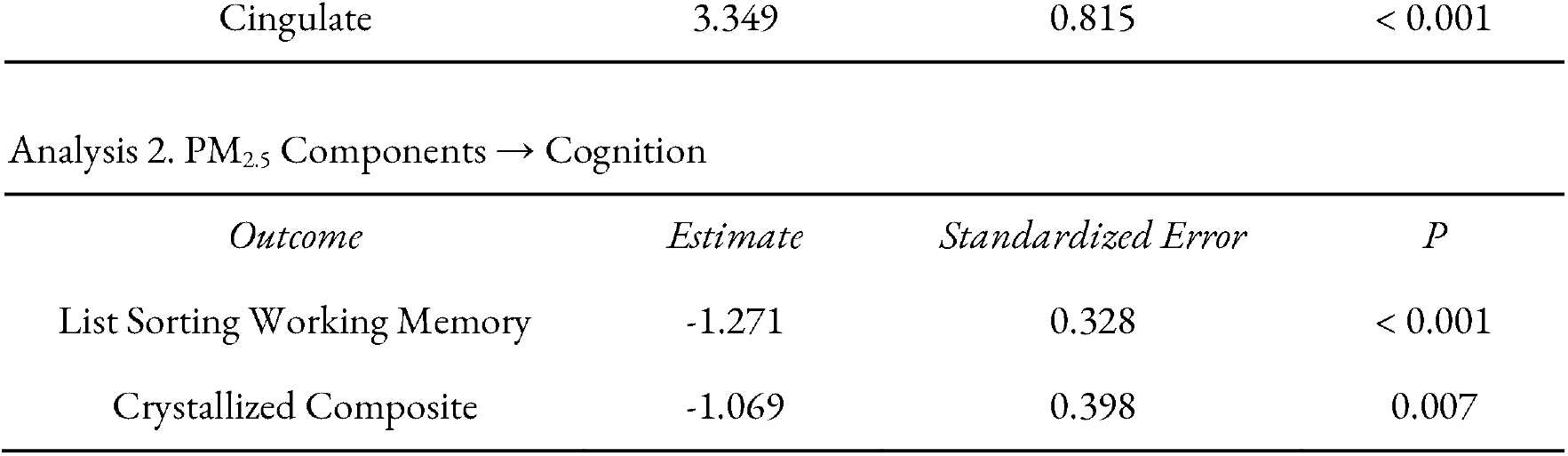
Results from multi-pollutant weighted quantile sum models (P < 0.01). Abbreviations: Perivascular Space (PVS).

#### 3.2.2 Analysis 2. Air Pollution and Cognition

WQS models examining associations between co-exposure and cognitive domains showed that co-exposure was associated with lower list sorting working memory (P < 0.001), where NH_4_ ^+^, NO _3_ ^-^, SO_4_ ^2-^, Zn, and Ca drove the association, and crystallized composite (P < 0.01), where NH _4_ ^+^, Si, Ca, and K drove the association (**Table 3, Figure S8, Table S8)**.

## 4. Discussion

### 4.1 Key Results

This study identified novel associations between air pollution exposure, PVS morphology, and cognition in preadolescents. Using a large, multi-ethnic sample of 9-10-year-olds from across the U.S., we identified distinct pollutant-PVS-cognition relationships. In single-pollutant (LME) models, NH_4_^+^ exposure was linked to greater temporal PVS VF and poorer language and crystallized intelligence, or an individual’s accumulation of knowledge; temporal PVS count was negatively associated with episodic and working memory, and fluid intelligence, or their ability to think flexibly. Zn was associated with higher frontal and CSO PVS count, which were in turn negatively associated with episodic memory and fluid cognition. Frontal count mediated the association between Zn and total cognition. Br was linked to higher cingulate PVS count which was associated with lower episodic and working memory, and total cognition. Ca exposure was associated with worse receptive language and processing speed. In contrast, Cu exposure was positively associated with expressive language and crystallized cognition. In multi-pollutant models, co-exposure to the 15 PM_2.5_ components was associated with higher temporal and cingulate PVS count, with Br, Cu, Si and NH_4_^+^ driving these relationships. Co-exposure was also associated with worse working memory and crystallized cognition, with NH_4_^+^, among others, driving these associations.

Negative associations were identified between PVS count and memory and all three cognitive summary scores. While associations with cognition were most commonly seen for the temporal and frontal lobes, and cingulate, significant associations existed for all examined regions except the occipital lobe (P_FDR_ < 0.01), suggesting global effects. These results are consistent with studies in adults, which have tied lower glymphatic system function and PVS burden to worse memory and overall cognitive function^19,20,56^. One potential explanation for the observed results is that higher PVS counts are indicative of disrupted waste clearance and neuroinflammation^57–59^, which may selectively harm brain regions involved in memory and executive function, such as the frontal and temporal lobes, and cingulate cortex.

Importantly, the associations in the current study are observed at low levels of air pollution exposure, highlighting that potential health risks are seen even below traditional regulatory thresholds. These results also suggest that the chemical composition of PM_2.5_ may be critical for determining neurovascular and cognitive effects, underscoring the need for air quality regulations that consider particle chemical composition rather than the mass concentration alone.

#### 4.1.1 Biological Mechanisms Linking Air Pollution Exposure to PVS Morphology

Considering the role of PVS in glymphatic function, waste clearance, and regulation of neuroinflammation^12^, and evidence that PVS and glymphatic dysfunction contribute to brain health and risk for disease^26,57,59^, pollution exposure may exacerbate systemic and neuroinflammation and increase metal load, thereby impairing glymphatic clearance and elevating risk for neurological disorders such as AD^25,26,60^. As PM_2.5_ particles cross the BBB and are dispersed throughout the brain, they can bind to metabolic waste, proteins, and neurotransmitters, thus upregulating neurotransmitter release, decreasing the solubility of waste, and inhibiting waste clearance^61^. For instance, exposure to NH_4_^+^ has been associated with changes in astrocyte and neuron morphology and apoptosis^62–64^. NH_4_^+^ can permeate epithelia via tight junctions^65^, where it can compete at K+ channel binding sites and alter neuronal excitability^66^It may also compromise astrocytic buffering by increasing extracellular K^+^ concentrations and over activating the Na^+^-K^+^-Cl^-^ cotransporter, which in turn depolarizes neuronal GABA and impairs cortical inhibitory networks^67^.

Metal exposure is particularly complex, as normal physiological functions depend on metal homeostasis and metals such as Zn and Cu are essential for human health and neuronal function^68,69^. However, metal exposure can have toxic effects both directly and through their effects on metal homeostasis. Metals in air pollution contribute to redox generative free radicals, oxidative stress in vivo, and metal dyshomeostasis^70^, which can then contribute to neurodevelopmental and neurodegenerative conditions^71^. For instance, Zn is directly tied to neuronal cell death at too high doses^72,73^ and can cause secondary toxicity via its interference with Cu uptake^74,75^. While Cu can interfere with long term potentiation^76,77^, it is also critical for normal cognitive function^78^. These complex relationships between various metals could be one reason that we see inverse relationships between Cu and Zn and cognition (i.e., Cu is positively associated with cognition while Zn is negatively associated), emphasizing the importance of metal homeostasis for proper neurological functioning and the capacity of air pollution to disrupt this balance.

Our results extend previous studies linking exposure to PM^2.5^ and its components to brain structure^79–81^ in pre- and early adolescents. Pollutant-brain associations identified by previous studies are distinct depending on the timing of exposure, methodological approach (i.e., single vs. multi-pollutant model), and pollutant or component assessed^1^. In the context of previous evidence, the present findings support the hypothesis that outdoor particle pollutants have differential neurotoxic potential, consistent with toxicological evidence from animal and cellular studies^71^.

#### 4.1.2 Broader Implications: A potential early marker of neurological disease risk?

PVS alterations in late childhood and early adolescence may represent an early biomarker of vascular or glymphatic dysfunction, with potential implications for long-term cognitive health. For instance, several PM_2.5_ components identified in the current study have also been linked to AD susceptibility and pathophysiology. Elevated ammonia (the neutral form of NH_4_^+^) can increase GABA release, upregulating inhibitory neurotransmission and contributing to cognitive deficits in AD^82^. Higher brain Br levels have been associated with AD neuropathology^83^. Accumulation of metals such as Fe, Cu, Ni, and Zn has been associated with Aβ and pTau aggregation^84^, and Fe, Cu, and Zn have been found in elevated concentrations in AD senile plaques^85,86^. Elevated serum calcium has been associated with faster cognitive decline in older adults^87^ and AD risk^88^. Finally, PVS burden and glymphatic system dysfunction–key for clearing interstitial solutes such as Aβ^89^–have been strongly associated with cognitive decline and AD^59,90^.

### 4.2 Strengths & Limitations

There are several strengths and limitations to note. First, the study provides novel insights into a highly understudied, yet critically important, research topic. The study utilizes a large, diverse sample of 9-10-year-olds from across the U.S., improving the generalizability of findings. Furthermore, the study leverages both single- and multi-pollutant models. While LMEs allow us to identify pollutant-specific exposure-brain-cognition relationships, WQS sheds light on the impact of co-exposures by minimizing the impact of collinearity between the correlated components while also identifying cumulative associations. However, due to image quality constraints, we were only able to retain data from a subset of ABCD Study participants. Compared to the whole ABCD Study, our participants were older and more likely to be female, white, and scanned on a Siemens scanner (**Table S1**), demonstrating a limitation for generalizability. Also, with regards to generalizability, participants in this sample are exposed to relatively low levels of air pollution, meaning that these results only provide insight into the impact of one year of low-level air pollution exposure and do not necessarily translate to children living in highly polluted countries. Finally, the current study utilizes cross-sectional design, limiting our ability to draw conclusions regarding the impacts of chronic exposure on neurodevelopmental and/or cognitive trajectories. Future studies should consider utilizing longitudinal models to answer these questions.

## 5. Conclusions

The current study identified novel associations between PM_2.5_ component exposure, PVS count, and cognition in preadolescents from across the United States. Results from this study highlight the brain’s susceptibility to the toxic effects of air pollution during the transition to adolescence and, for the first time, suggest that PVS dysfunction may be one mechanism by which air pollution harms cognitive functioning in preadolescents.

## Acknowledgments

A special thank you to all participants and their families for their participation in the ABCD Study. This work was supported by the National Institutes of Health (NIH) National Institute of Environmental Health Sciences (NIEHS) (Grant Nos. R01ES032295 and R01ES031074 [to MMH]; T32ES013678 [to JM]; P30ES07048 [to JM and MAR]; 3P30ES000002-55S [to MAR]), National Institute of Mental Health (NIMH) (Grant RF1MH123223 [to JC]), National Institute of Neurological Disorders and Stroke (Grant R01NS128486 [to JC]), and EPA grants (Grant Nos. 83587201 and 83544101 [to JS]).

Data used in the preparation of this article were obtained from the Adolescent Brain Cognitive Development^SM^ (ABCD) Study (https://abcdstudy.org), held in the NIMH Data Archive (NDA). This is a multisite, longitudinal study designed to recruit more than 10,000 children aged 9-10 and follow them over 10 years into early adulthood. The ABCD Study® is supported by the National Institutes of Health and additional federal partners under award numbers U01DA041048, U01DA050989, U01DA051016, U01DA041022, U01DA051018, U01DA051037, U01DA050987, U01DA041174, U01DA041106, U01DA041117, U01DA041028, U01DA041134, U01DA050988, U01DA051039, U01DA041156, U01DA041025, U01DA041120, U01DA051038, U01DA041148, U01DA041093, U01DA041089, U24DA041123, U24DA041147. A full list of supporters is available at https://abcdstudy.org/federal-partners.html. A listing of participating sites and a complete listing of the study investigators can be found at https://abcdstudy.org/consortium_members/. ABCD consortium investigators designed and implemented the study and/or provided data but did not necessarily participate in the analysis or writing of this report. This manuscript reflects the views of the authors and may not reflect the opinions or views of the NIH or ABCD consortium investigators. The ABCD data repository grows and changes over time. The ABCD data used in this report came from DOI: 10.15154/8873-zj65 and DOI: 10.15154/1520591.

## Declaration of Interests

The authors declare no competing interests. Data Sharing

Data used in the preparation of this article were obtained from the Adolescent Brain Cognitive Development (ABCD) Study (https://abcdstudy.org), held in the NIMH Data Archive (NDA), releases 3.0 (DOI: 10.15154/8873-zj65) and 5.0 (DOI: 10.15154/1520591). Individual participant data will not be made available.

## Author Contributions

Conceptualization: JM, MMH

Methodology: JM

Formal analysis: JM, KS

Resources: MMH, JS, JC

Data curation: JM, CT, HL

Writing - original draft: JM

Writing - review and editing: JM, KS, MAR, MMH, JC

Visualization: JM

Supervision: MMH, JC

Project administration: MMH

Funding acquisition: JM, MMH, JC, JS, J-CC

