## Supplement for "Outdoor Air Pollution, Perivascular Space Morphology, and Cognition in Preadolescence"

**Supplemental Material for Outdoor Air Pollution, Perivascular Space Morphology, and Cognition in Preadolescence**

### **A. Supplemental Methods**

#### **A.1 Neuroimaging**

A harmonized data protocol was utilized across all ABCD study sites^1^. To reduce motion distortion, motion compliance training and real-time, prospective motion correction were utilized. T1-weighted (T1w) images were acquired using a magnetization-prepared rapid acquisition gradient echo sequence; T2-weighted (T2w) images were obtained with a fast spin echo sequence with variable flip angle^1^. T1w and T2w acquisitions consist of 176 slices with 1 mm^3^ isotropic resolution. Images were then reviewed by study staff and only data meeting quality control standards for recommended inclusion and those without clinical findings^2^ were included.

The current study uses our recently developed PVS segmentation pipeline^3^, which uses Frangi filter 3-dimensional vesselness tubular structure probability estimation as a salient estimation to train a Recurrent Neural Network (RNN) for mapping PVS morphology. First, raw T1w and T2w MRI images were registered using the Human Connectome Project (HCP) minimal preprocessing pipeline^4^. Next, adaptive non-local mean filtering of high-frequency spatial noise is applied to the preprocessed images to preserve PVS voxels while still removing noise. To do so, filtering is applied on high frequency noise using a one voxel frequency patch^5,6^. Next, an enhanced PVS contrast (EPC) image is generated by dividing the preprocessed T1w image by the preprocessed T2w image, which increases the visibility of white matter PVS contrast^5^. The WPSS pipeline is then applied, which utilizes the Frangi Filter^7^ to direct the segmentation process to focus on highlighting key areas corresponding to tubular structures, in this case PVS. While traditional fully supervised techniques required pixel-level annotations, WPSS uses weak supervision by incorporating key features extracted from the Frangi filter, which allows the model to learn from fewer labeled data, reducing the need for extensive manual annotations while maintaining high segmentation accuracy. Then, a U-Net convolutional neural network^8^ is trained to distinguish PVS from surrounding tissues, followed by implementation of a Conditional Random Field (CRF) as an RNN module^9^ to refine the segmentation by utilizing the salient guidance from the Frangi filter. This two-step method ensures precise segmentation even with limited annotated data. The WPSS model weights used in the current study was trained on the HCP dataset^5,10,11^, using 400 EPC images generated by the pipeline described by Sepehrband et al. (2019). Please see Lan et al. (2023) for additional details.

#### **A.2 Weighted Quantile Sum (WQS) Regression Methods**

WQS was executed as previously published by our team^12^. First, the mixture components are converted into quantile scores such that extreme values of component exposures do not overpower the weight estimation. The input data is then randomly split into training (40%) and validation (60%) subset samples as per common practice using the *gwqs* R package^13^. Then to construct the index, the training data is used to fit a generalized regression model using the quantile exposure variables and relevant covariates. Weights for each PM component are derived using maximum likelihood estimation procedure across 1000 bootstrap iterations, with the constraint that all weights sum to one. These weights are averaged across all bootstraps to create a single WQS index that integrates all 15 PM_2.5_ component exposures. Next, this constructed WQS index is used to estimate the overall mixture effect on the outcome using the remaining validation data to quantify the association (coefficients) and determine the statistical significance (p-values).

#### **A.3 Analyses**

Five components—EC, NH_4_, NO_3_, OC, SO_4_—were converted from ug/m^3^ to ng/m^3^ to match the units of the other 10 components.

##### A.3.1 Analysis 1: Air Pollution and Perivascular Space (PVS) Morphology

To first answer the question of whether air pollution exposure was associated with PVS morphology (i.e., count and volume), we ran two sets of single-pollutant linear mixed-effects (LME) models using the *lme4::lmer()* function in R. First, we explored how each of the 18 pollutants associated with PVS count in six regions (frontal, temporal, parietal, occipital, centrum semiovale (CSO), and cingulate) using the following formula:

*Regional PVS Count ~ 1 + Pollutant + Age + Sex + Race/Ethnicity + Household Income + BMI-z + Scanner/Coil + Regional Volume + (1 | Site)*

Then, we explored how each of the 18 pollutants associated with PVS volume fraction in six regions using the same formula as above, but dropping regional volume as a covariate as the outcome already accounts for differences in cranial size by dividing regional PVS volume / regional volume.

*Regional PVS Volume Fraction ~ 1 + Pollutant + Age + Sex + Race/Ethnicity + Household Income + BMI-z + Scanner/Coil + (1 | Site)*

##### A.3.2 Analysis 2: PVS Morphology and Cognition

Our next set of analysis was aimed at exploring the functional implications of differences in PVS morphology in 9-10-year-olds. Considering that our initial analyses mainly identified significant relationships between air pollution exposure and PVS count, not volume fraction, we made an a priori decision to only perform post-hoc analyses using count as the main predictor. Therefore, we performed a set of LME models testing how PVS count in each of the six regions related to age-corrected scores across 10 NIH Toolbox Cognitive domains. The formula used is as follows:

*NIH TB Age-Adjusted Score ~ 1 + PVS Regional Count + Sex + Race/Ethnicity + Household Income + BMI-z + Scanner/Coil + Regional Volume + (1 | Site)*

##### A.3.3 Analysis 3: Air Pollution and Cognition

To test whether exposure to air pollution was associated with cognition in 9-10-year-olds, we use LME models to test whether exposure to 18 pollutants was associated with 10 NIH Toolbox Cognitive outcomes. The formula used is as follows:

*NIH TB Age-Adjusted Score ~ 1 + Pollutant + Sex + Race/Ethnicity + Household Income + BMI-z + Scanner/Coil + (1 | Site)*

False discovery rate (FDR) correction was implemented following each set of analyses described above (**A.3.1-A.3.3**) using the *stats::p.adjust*() function in *R* and only results passing FDR correction for significance at P_FDR_ < 0.01 were considered for mediation analysis. Cohen’s effect size was calculated using the *effectsize::cohens_f_squared*() function.

##### A.3.4 Analysis 4: Mediation Analysis

After running Analyses 1-3, we conducted one mediation analysis using the *mediation* package in R and the Baron and Kenny method^14^ to assess whether PVS count mediates the relationship between air pollution exposure and cognition. To do so, we ran the following:

*NIH TB Age-Adjusted Total Composite ~ 1 + Frontal PVS Count + Zinc + Age + Sex + Race/Ethnicity + Household Income + BMI-z + Scanner/Coil + CSO Volume + (1 | Site)*

##### A.3.5 Analysis 4: Weighted Quantile Sum (WQS) Regression

The generalized WQS regression expression takes the following form^15^:

$g(\mu) = {\beta_{0} +\beta_{1}}\left( \sum_{i = 1}^{15} w_{i}q_{i} \right) + z'\varphi$

Where g is a differentiable link function(monotonic) that relates the mean, μ, with the independent variables on the right hand side of the equation. i iterates through the 15 mixture components of PM_2.5_. The weight for the i^th^ component concentration is given by w_i_ conditional such that the sum of these weights equal to 1 and the quantile score for that component concentration is q_i,_ for a given observation. The covariates adjusted for in the model are denoted by z. By averaging across the weights from each of the 1000 bootstraps, the WQS model generates an overall mixture index which is a composite index inclusive of all the 15 PM_2.5_ component exposures, represented as $\sum_{i = 1}^{15} w_{i}q_{i}$.

### **B. Tables**

#### **Table S1. Performance of air pollution models.** Abbreviations: Fine Particulate Matter (PM_2.5_), Nitrogen Dioxide (NO_2_), Ozone (O_3_); Bromine (Br), Calcium (Ca), Copper (Cu), Elemental Carbon (EC), Iron (Fe), Potassium (K), Ammonium (NH_4_^+^), Nickel (Ni), Nitrate (NO_3_^-^), Organic Carbon (OC), Lead (Pb), Silicon (Si), Sulfate (SO_4_^2-^), Vanadium (V), Zinc (Zn); Cross Validated (CV).

| **Pollutant** | **CV (R^2^)** |
| --- | --- |
| *Criteria Pollutant* |  |
| PM_2.5_ | 0.86 |
| NO_2_ | 0.79 |
| O_3_ | 0.86 |
| *PM_2.5_ Component* |  |
| Br | 0.86 |
| Ca | 0.86 |
| Cu | 0.79 |
| EC | 0.91 |
| Fe | 0.87 |
| K | 0.87 |
| NH_4_^+^ | 0.91 |
| Ni | 0.85 |
| NO_3_^-^ | 0.88 |
| OC | 0.86 |
| Pb | 0.83 |
| Si | 0.85 |
| SO_4_^2-^ | 0.95 |
| V | 0.84 |
| Zn | 0.88 |

#### **Table S2. Sample characteristics of the whole ABCD Study cohort and final study sample.** Total number (N) of participants and (percentage of total sample) for each demographic category are presented for the whole ABCD cohort and the final study sample. Abbreviations: Adolescent Brain Cognitive Development (ABCD) Study; General Educational Development (GED); General Electric (GE); Body Mass Index (BMI).

|  | **Whole ABCD Cohort** | **Study Sample** |
| --- | --- | --- |
|  | **N (%)** | **N (%)** |
| Total N | 11,832 (100%) | 6,949 (58.7%) |
| Sex (F)*** | 5,655 (47.8%) | 3,414 (49.1%) |
| *Race & Ethnicity*** |  |  |
| Asian | 250 (2.1%) | 161 (2.3%) |
| Hispanic | 2,404 (20.3%) | 1,456 (21.0%) |
| Black | 1,777 (15.0%) | 920 (13.2%) |
| White | 6,156 (52.1%) | 3,720 (53.5%) |
| Other | 1,243 (10.5%) | 691 (9.9%) |
| *Household Income* |  |  |
| < $50,000 | 3,214 (27.2%) | 1,779 (25.6%) |
| ≥$50,000 to <$100,000 | 3,063 (25.9%) | 1,846 (26.6%) |
| ≥ $100,000 | 4,541 (38.4%) | 2,746 (39.5%) |
| Don’t Know / Refuse | 1,012 (8.6%) | 578 (8.3%) |
| *Caregiver Education* |  |  |
| < High school diploma | 592 (5.0%) | 318 (4.6%) |
| High school diploma / GED | 1,129 (9.5%) | 632 (9.1%) |
| Some college / Associate’s degree | 3,068 (26.0%) | 1,737 (25.0%) |
| Bachelor’s degree | 3,005 (25.4%) | 1,776 (25.6%) |
| Graduate degree | 4,024 (34.0%) | 2,481 (35.7%) |
| *MRI Scanner Manufacturer**** |  |  |
| Siemens | 7,273 (62.0%) | 4,705 (67.7%) |
| GE Medical Systems | 2,939 (25.0%) | 1,429 (20.6%) |
| Philips Medical Systems | 1,523 (13.0%) | 815 (11.7%) |
|  | **Mean (SD)** | **Mean (SD)** |
| Age at baseline (months)* | 119.0 (7.5) | 119.2 (7.5) |
| BMI Z-Score | 0.4 (1.2) | 0.4 (1.2) |

*Note.* Differences in sample characteristics between the study sample and whole ABCD sample have been tested using Pearson’s Chi-Square or t-tests, as appropriate. Stars in column one indicate test significance, where * = p < 0.05, ** = p < 0.01, and *** = p < 0.001. Items without stars (i.e., Household Income) were not statistically significantly different between samples. “Other” Race/Ethnicity category includes participants identified by their caregiver as American Indian/Native American, Alaska Native, Native Hawaiian, Guamanian, Samoan, Other Pacific Islander, Asian Indian, Chinese, Filipino, Japanese, Korean, Vietnamese, Other Asian not listed, or Other Race not listed.

#### **Table S3. Descriptive statistics for participant exposures to annual average criteria pollutants and fine particulate matter (PM_2.5_) components.** Mean and standard deviation (SD) are presented for each criteria pollutant and PM_2.5_ component. Exposures included for the whole ABCD Study cohort and the final study sample. Abbreviations: Fine Particulate Matter (PM_2.5_), Nitrogen Dioxide (NO_2_), Ozone (O_3_); Bromine (Br), Calcium (Ca), Copper (Cu), Elemental Carbon (EC), Iron (Fe), Potassium (K), Ammonium (NH_4_^+^), Nickel (Ni), Nitrate (NO_3_^-^), Organic Carbon (OC), Lead (Pb), Silicon (Si), Sulfate (SO_4_^2-^), Vanadium (V), Zinc (Zn); Parts per Billion (ppb).

|  | **Whole ABCD Cohort Mean (SD)** | **Study Sample**  **Mean (SD)** |
| --- | --- | --- |
| Total N | 11,832 | 6,949 |
| *Criteria Pollutants* |  |  |
| PM_2.5_ (*µg/m^3^*) | 7.7 (1.6) | 7.6 (1.6) |
| NO_2_ (*ppb*) | 18.6 (5.8) | 18.7 (5.8) |
| O_3_ (*ppb*)*** | 41.5 (4.4) | 41.8 (4.5) |
| *PM_2.5_ Components (ng/m^3^)* |  |  |
| Br | 2.6 (0.6) | 2.6 (0.6) |
| Ca | 48.1 (20.9) | 48.1 (21.3) |
| Cu | 4.6 (1.7) | 4.6 (1.7) |
| EC | 523.2 (159.2) | 526.8 (162.9) |
| Fe | 65.5 (24.7) | 65.8 (25.3) |
| K | 62.7 (9.8) | 62.9 (9.9) |
| NH_4_^+^ *** | 301.4 (122.5) | 293.9 (124.3) |
| Ni | 0.8 (0.3) | 0.8 (0.3) |
| NO_3_^-^ * | 927.7 (349.3) | 914.7 (360.8) |
| OC | 1,884.1 (453.6) | 1,887.8 (468.6) |
| Pb* | 4.5 (1.2) | 4.5 (1.3) |
| Si | 83.7 (37.9) | 84.4 (38.3) |
| SO_4_^2-^ * | 919.4 (286.3) | 908.8 (292.8) |
| V | 0.4 (0.2) | 0.4 (0.2) |
| Zn | 9.3 (4.1) | 9.2 (4.1) |

*Note.* Differences in sample characteristics between the study sample and whole ABCD sample have been tested using t-tests. Stars in column one indicate test significance, where * = p < 0.05, ** = p < 0.01, and *** = p < 0.001. Items without stars (i.e., PM_2.5_) were not statistically significantly different between samples.

#### **Table S4. Descriptive statistics for PVS counts and volume fraction (mm^3^) at baseline (9-10 years) in the final study sample from the ABCD Study cohort.** Mean and standard deviation (SD) are presented for PVS count and volume fraction (mm^3^) in each key brain region. Abbreviations: Perivascular Space (PVS).

|  | **Mean (SD)** |
| --- | --- |
| Total N | 6,949 |
| *PVS Count* |  |
| Frontal Lobe | 832.9 (267.5) |
| Temporal Lobe | 241.8 (69.6) |
| Parietal Lobe | 448.4 (139.2) |
| Occipital Lobe | 124.2 (41.3) |
| Centrum Semiovale | 1,029.7 (323.3) |
| Cingulate | 189.7 (63.0) |
| *PVS Volume Fraction (mm^3^)* |  |
| Frontal Lobe | 0.006 (0.002) |
| Temporal Lobe | 0.003 (0.001) |
| Parietal Lobe | 0.005 (0.002) |
| Occipital Lobe | 0.002 (0.001) |
| Centrum Semiovale | 0.006 (0.002) |
| Cingulate | 0.005 (0.002) |

**Table S5. Results from single-pollutant linear mixed-effects models examining relationships between air pollution exposure and PVS count and volume fraction.** * = P_FDR_ < 0.05; ** = P_FDR_ < 0.01; *** = P_FDR_ < 0.001. Abbreviations: Standardized (Std.), Confidence Interval (CI); Fine Particulate Matter (PM_2.5_), Nitrogen Dioxide (NO_2_), Ozone (O_3_), Bromine (Br), Calcium (Ca), Copper (Cu), Elemental Carbon (EC), Iron (Fe), Potassium (K), Ammonium (NH_4_^+^), Nitrate (NO_3_^-^), Nickel (Ni), Organic Carbon (OC), Lead (Pb), Sulfate (SO_4_^2-^), Silicon (Si), Vanadium (V), Zinc (Zn); False Discovery Rate (FDR); Centrum Semiovale (CSO); Volume Fraction (VF).

| **Temporal VF** | | | | | **Temporal Count** | | | |
| --- | --- | --- | --- | --- | --- | --- | --- | --- |
| ***Pollutant*** | ***Std. Beta*** | ***CI*** | ***P_FDR_*** | ***Cohen’s F^2^*** | ***Std. Beta*** | ***CI*** | ***P_FDR_*** | ***Cohen’s F^2^*** |
| **NO_2_** | 0.025 | -0.006-0.056 | 0.201 | 0.017 | 0.012 | -0.015-0.039 | 0.485 | 0.002 |
| **O_3_** | -0.005 | -0.033-0.022 | 0.803 | 0.001 | -0.003 | -0.025-0.02 | 0.812 | < 0.001 |
| **PM_2.5_** | 0.038 | 0.006-0.07 | 0.057 | 0.034 | 0.025 | -0.003-0.053 | 0.173 | 0.005 |
| **Br** | 0.021 | -0.014-0.056 | 0.31 | 0.006 | 0.045 | 0.013-0.077 | 0.02* | 0.008 |
| **Ca** | 0.033 | -0.009-0.075 | 0.202 | 0.005 | 0.026 | -0.012-0.064 | 0.327 | 0.001 |
| **Cu** | 0.037 | 0.006-0.068 | 0.057 | 0.04 | 0.035 | 0.009-0.061 | 0.025* | 0.015 |
| **EC** | 0.024 | -0.008-0.056 | 0.202 | 0.013 | 0.018 | -0.01-0.045 | 0.33 | 0.003 |
| **Fe** | 0.054 | 0.02-0.089 | 0.02* | 0.045 | 0.047 | 0.016-0.077 | 0.016* | 0.011 |
| **K** | 0.038 | 0.005-0.071 | 0.06 | 0.029 | 0.042 | 0.014-0.07 | 0.017* | 0.013 |
| **NH_4_^+^** | 0.065 | 0.03-0.1 | 0.009** | 0.053 | 0.059 | 0.023-0.094 | 0.016* | 0.007 |
| **Ni** | 0.015 | -0.013-0.043 | 0.346 | 0.012 | 0.011 | -0.012-0.033 | 0.485 | 0.003 |
| **NO_3_^-^** | 0.054 | 0.015-0.092 | 0.032* | 0.022 | 0.05 | 0.012-0.089 | 0.026* | 0.003 |
| **OC** | 0.029 | -0.01-0.067 | 0.202 | 0.006 | 0.017 | -0.019-0.053 | 0.485 | 0.001 |
| **Pb** | 0.031 | -0.002-0.064 | 0.131 | 0.018 | 0.022 | -0.005-0.05 | 0.23 | 0.004 |
| **Si** | 0.005 | -0.039-0.049 | 0.81 | < 0.001 | 0.017 | -0.026-0.059 | 0.535 | < 0.001 |
| **SO_4_^2-^** | 0.005 | -0.039-0.049 | 0.81 | < 0.001 | 0.005 | -0.038-0.048 | 0.812 | < 0.001 |
| **V** | -0.032 | -0.067-0.002 | 0.131 | 0.015 | -0.009 | -0.039-0.021 | 0.628 | < 0.001 |
| **Zn** | 0.045 | 0.013-0.077 | 0.032* | 0.043 | 0.045 | 0.016-0.073 | 0.016* | 0.015 |
| **Parietal VF** | | | | |  | **Parietal Count** | | |
| ***Pollutant*** | ***Std. Beta*** | ***CI*** | ***P_FDR_*** | ***Cohen’s F^2^*** | ***Std. Beta*** | ***CI*** | ***P_FDR_*** | ***Cohen’s F^2^*** |
| **NO_2_** | 0.015 | -0.013-0.043 | 0.894 | 0.009 | 0.026 | 0.001-0.052 | 0.106 | 0.015 |
| **O_3_** | -0.007 | -0.033-0.018 | 0.894 | 0.004 | -0.003 | -0.025-0.019 | 0.847 | < 0.001 |
| **PM_2.5_** | 0.006 | -0.023-0.035 | 0.894 | 0.001 | 0.018 | -0.008-0.045 | 0.287 | 0.006 |
| **Br** | -0.006 | -0.036-0.024 | 0.894 | 0.001 | 0.017 | -0.013-0.046 | 0.403 | 0.003 |
| **Ca** | 0.009 | -0.027-0.045 | 0.894 | 0.001 | 0.008 | -0.027-0.044 | 0.718 | < 0.001 |
| **Cu** | -0.005 | -0.033-0.023 | 0.894 | 0.001 | 0.02 | -0.005-0.045 | 0.221 | 0.01 |
| **EC** | -0.003 | -0.031-0.026 | 0.899 | < 0.001 | 0.027 | 0-0.053 | 0.106 | 0.012 |
| **Fe** | 0.019 | -0.011-0.05 | 0.894 | 0.011 | 0.042 | 0.013-0.071 | 0.027* | 0.019 |
| **K** | 0.009 | -0.02-0.038 | 0.894 | 0.003 | 0.03 | 0.003-0.057 | 0.097 | 0.013 |
| **NH_4_^+^** | 0.029 | -0.003-0.061 | 0.894 | 0.016 | 0.05 | 0.019-0.081 | 0.018* | 0.015 |
| **Ni** | -0.006 | -0.032-0.021 | 0.894 | 0.002 | 0.018 | -0.005-0.04 | 0.224 | 0.014 |
| **NO_3_^-^** | 0.023 | -0.01-0.057 | 0.894 | 0.008 | 0.039 | 0.005-0.074 | 0.094 | 0.005 |
| **OC** | 0.004 | -0.029-0.037 | 0.894 | < 0.001 | 0.016 | -0.017-0.049 | 0.455 | 0.001 |
| **Pb** | 0.007 | -0.024-0.038 | 0.894 | 0.001 | 0.032 | 0.005-0.058 | 0.088 | 0.015 |
| **Si** | 0.006 | -0.03-0.042 | 0.894 | < 0.001 | -0.002 | -0.039-0.036 | 0.933 | < 0.001 |
| **SO_4_^2-^** | -0.006 | -0.042-0.03 | 0.894 | < 0.001 | 0.015 | -0.023-0.052 | 0.556 | 0.001 |
| **V** | -0.018 | -0.05-0.013 | 0.894 | 0.008 | -0.007 | -0.036-0.021 | 0.718 | 0.001 |
| **Zn** | -0.002 | -0.031-0.028 | 0.908 | < 0.001 | 0.044 | 0.017-0.071 | 0.018* | 0.029 |
| **Occipital VF** | | | | | **Occipital Count** | | | |
| ***Pollutant*** | ***Std. Beta*** | ***CI*** | ***P_FDR_*** | ***Cohen’s F^2^*** | ***Std. Beta*** | ***CI*** | ***P_FDR_*** | ***Cohen’s F^2^*** |
| **NO_2_** | 0.004 | -0.026-0.035 | 0.886 | < 0.001 | 0.009 | -0.016-0.035 | 0.985 | 0.002 |
| **O_3_** | -0.004 | -0.03-0.023 | 0.886 | 0.001 | -0.001 | -0.023-0.021 | 0.985 | < 0.001 |
| **PM_2.5_** | 0.009 | -0.023-0.04 | 0.763 | 0.001 | 0.006 | -0.021-0.032 | 0.985 | 0.001 |
| **Br** | -0.012 | -0.045-0.021 | 0.758 | 0.002 | -0.007 | -0.036-0.022 | 0.985 | 0.001 |
| **Ca** | 0.016 | -0.025-0.057 | 0.758 | 0.001 | 0.005 | -0.03-0.04 | 0.985 | < 0.001 |
| **Cu** | 0.011 | -0.019-0.041 | 0.758 | 0.003 | 0.019 | -0.007-0.044 | 0.664 | 0.008 |
| **EC** | -0.016 | -0.047-0.014 | 0.758 | 0.005 | < 0.001 | -0.026-0.026 | 0.985 | < 0.001 |
| **Fe** | 0.028 | -0.006-0.063 | 0.69 | 0.008 | 0.031 | 0.002-0.06 | 0.34 | 0.01 |
| **K** | 0.015 | -0.017-0.047 | 0.758 | 0.003 | 0.018 | -0.01-0.045 | 0.734 | 0.005 |
| **NH_4_^+^** | 0.031 | -0.008-0.07 | 0.69 | 0.005 | 0.03 | -0.003-0.063 | 0.467 | 0.004 |
| **Ni** | -0.001 | -0.028-0.027 | 0.966 | < 0.001 | 0.003 | -0.019-0.026 | 0.985 | < 0.001 |
| **NO_3_^-^** | 0.004 | -0.037-0.044 | 0.909 | < 0.001 | -0.004 | -0.039-0.031 | 0.985 | < 0.001 |
| **OC** | -0.017 | -0.053-0.019 | 0.758 | 0.002 | -0.002 | -0.034-0.03 | 0.985 | < 0.001 |
| **Pb** | 0.018 | -0.014-0.05 | 0.758 | 0.005 | 0.015 | -0.012-0.042 | 0.812 | 0.003 |
| **Si** | -0.012 | -0.054-0.03 | 0.763 | < 0.001 | -0.012 | -0.049-0.025 | 0.985 | < 0.001 |
| **SO_4_^2-^** | -0.013 | -0.056-0.029 | 0.763 | 0.001 | < 0.001 | -0.038-0.037 | 0.985 | < 0.001 |
| **V** | -0.018 | -0.051-0.016 | 0.758 | 0.003 | -0.007 | -0.035-0.021 | 0.985 | 0.001 |
| **Zn** | 0.026 | -0.006-0.058 | 0.69 | 0.01 | 0.035 | 0.008-0.062 | 0.199 | 0.018 |
| **Frontal VF** | | | | | **Frontal Count** | | | |
| ***Pollutant*** | ***Std. Beta*** | ***CI*** | ***P_FDR_*** | ***Cohen’s F^2^*** | ***Std. Beta*** | ***CI*** | ***P_FDR_*** | ***Cohen’s F^2^*** |
| **NO_2_** | 0.019 | -0.009-0.048 | 0.462 | 0.006 | 0.031 | 0.005-0.057 | 0.068 | 0.009 |
| **O_3_** | -0.008 | -0.033-0.016 | 0.703 | 0.003 | -0.003 | -0.025-0.019 | 0.783 | < 0.001 |
| **PM_2.5_** | 0.013 | -0.017-0.042 | 0.703 | 0.002 | 0.024 | -0.003-0.052 | 0.126 | 0.004 |
| **Br** | 0.008 | -0.025-0.04 | 0.703 | < 0.001 | 0.035 | 0.004-0.066 | 0.068 | 0.004 |
| **Ca** | 0.02 | -0.019-0.059 | 0.61 | 0.001 | 0.023 | -0.014-0.061 | 0.304 | 0.001 |
| **Cu** | 0.011 | -0.017-0.038 | 0.703 | 0.002 | 0.029 | 0.004-0.055 | 0.068 | 0.009 |
| **EC** | 0.005 | -0.024-0.034 | 0.727 | < 0.001 | 0.024 | -0.002-0.051 | 0.126 | 0.004 |
| **Fe** | 0.023 | -0.008-0.055 | 0.438 | 0.005 | 0.037 | 0.007-0.066 | 0.068 | 0.005 |
| **K** | 0.023 | -0.007-0.053 | 0.438 | 0.006 | 0.037 | 0.009-0.064 | 0.059 | 0.008 |
| **NH_4_^+^** | 0.042 | 0.007-0.078 | 0.377 | 0.007 | 0.046 | 0.011-0.081 | 0.059 | 0.003 |
| **Ni** | 0.006 | -0.019-0.03 | 0.703 | 0.001 | 0.017 | -0.005-0.039 | 0.186 | 0.007 |
| **NO_3_^-^** | 0.034 | -0.004-0.073 | 0.438 | 0.003 | 0.042 | 0.004-0.08 | 0.07 | 0.002 |
| **OC** | 0.01 | -0.026-0.046 | 0.703 | < 0.001 | 0.015 | -0.02-0.051 | 0.457 | < 0.001 |
| **Pb** | 0.018 | -0.011-0.048 | 0.5 | 0.004 | 0.028 | 0.001-0.054 | 0.082 | 0.006 |
| **Si** | 0.009 | -0.032-0.051 | 0.703 | < 0.001 | 0.018 | -0.024-0.06 | 0.457 | < 0.001 |
| **SO_4_^2-^** | -0.01 | -0.052-0.033 | 0.703 | < 0.001 | 0.013 | -0.029-0.055 | 0.579 | < 0.001 |
| **V** | -0.028 | -0.059-0.004 | 0.438 | 0.007 | -0.014 | -0.043-0.015 | 0.441 | 0.001 |
| **Zn** | 0.024 | -0.006-0.054 | 0.438 | 0.007 | 0.051 | 0.023-0.078 | 0.005** | 0.016 |
| **CSO VF** | | | | | **CSO Count** | | | |
| ***Pollutant*** | ***Std. Beta*** | ***CI*** | ***P_FDR_*** | ***Cohen’s F^2^*** | ***Std. Beta*** | ***CI*** | ***P_FDR_*** | ***Cohen’s F^2^*** |
| **NO_2_** | 0.016 | -0.012-0.044 | 0.76 | 0.007 | 0.028 | 0.003-0.054 | 0.074 | 0.01 |
| **O_3_** | -0.007 | -0.032-0.017 | 0.914 | 0.003 | -0.002 | -0.024-0.019 | 0.823 | < 0.001 |
| **PM_2.5_** | 0.009 | -0.02-0.038 | 0.914 | 0.002 | 0.022 | -0.004-0.049 | 0.168 | 0.005 |
| **Br** | 0.001 | -0.03-0.033 | 0.991 | < 0.001 | 0.029 | -0.001-0.06 | 0.117 | 0.004 |
| **Ca** | 0.013 | -0.025-0.051 | 0.914 | 0.001 | 0.017 | -0.02-0.053 | 0.468 | < 0.001 |
| **Cu** | 0.004 | -0.024-0.032 | 0.991 | < 0.001 | 0.027 | 0.002-0.052 | 0.074 | 0.011 |
| **EC** | < 0.001 | -0.029-0.029 | 0.991 | < 0.001 | 0.024 | -0.002-0.051 | 0.129 | 0.006 |
| **Fe** | 0.019 | -0.012-0.05 | 0.76 | 0.006 | 0.038 | 0.009-0.068 | 0.049 | 0.009 |
| **K** | 0.017 | -0.013-0.047 | 0.76 | 0.006 | 0.035 | 0.008-0.063 | 0.049 | 0.011 |
| **NH_4_^+^** | 0.036 | 0.002-0.07 | 0.682 | 0.011 | 0.049 | 0.015-0.082 | 0.045 | 0.006 |
| **Ni** | 0.001 | -0.024-0.026 | 0.991 | < 0.001 | 0.018 | -0.004-0.04 | 0.173 | 0.01 |
| **NO_3_^-^** | 0.028 | -0.008-0.064 | 0.751 | 0.005 | 0.04 | 0.003-0.077 | 0.074 | 0.002 |
| **OC** | 0.007 | -0.027-0.042 | 0.936 | < 0.001 | 0.016 | -0.019-0.05 | 0.468 | 0.001 |
| **Pb** | 0.016 | -0.014-0.046 | 0.76 | 0.005 | 0.032 | 0.006-0.059 | 0.063 | 0.01 |
| **Si** | 0.002 | -0.038-0.041 | 0.991 | < 0.001 | 0.009 | -0.032-0.05 | 0.698 | < 0.001 |
| **SO_4_^2-^** | -0.01 | -0.049-0.03 | 0.936 | < 0.001 | 0.014 | -0.027-0.054 | 0.569 | < 0.001 |
| **V** | -0.026 | -0.057-0.005 | 0.751 | 0.01 | -0.012 | -0.041-0.017 | 0.488 | 0.001 |
| **Zn** | 0.014 | -0.016-0.044 | 0.792 | 0.004 | 0.05 | 0.023-0.077 | 0.005 | 0.022 |
| **Cingulate VF** | | | | | **Cingulate Count** | | | |
| ***Pollutant*** | ***Std. Beta*** | ***CI*** | ***P_FDR_*** | ***Cohen’s F^2^*** | ***Std. Beta*** | ***CI*** | ***P_FDR_*** | ***Cohen’s F^2^*** |
| **NO_2_** | 0.028 | -0.003-0.059 | 0.28 | 0.002 | 0.027 | 0-0.054 | 0.22 | 0.001 |
| **O_3_** | -0.014 | -0.039-0.012 | 0.544 | 0.002 | -0.012 | -0.035-0.01 | 0.498 | 0.001 |
| **PM_2.5_** | 0.015 | -0.018-0.047 | 0.573 | < 0.001 | 0.015 | -0.014-0.044 | 0.498 | < 0.001 |
| **Br** | 0.055 | 0.017-0.093 | 0.078 | 0.003 | 0.082 | 0.048-0.116 | < 0.001 | 0.003 |
| **Ca** | 0.025 | -0.021-0.07 | 0.544 | < 0.001 | 0.015 | -0.026-0.056 | 0.607 | < 0.001 |
| **Cu** | 0.031 | 0.001-0.061 | 0.234 | 0.004 | 0.039 | 0.012-0.065 | 0.037 | 0.004 |
| **EC** | < 0.001 | -0.032-0.032 | 0.986 | < 0.001 | < 0.001 | -0.029-0.028 | 0.973 | < 0.001 |
| **Fe** | 0.026 | -0.01-0.061 | 0.354 | 0.001 | 0.024 | -0.007-0.056 | 0.325 | < 0.001 |
| **K** | 0.043 | 0.01-0.076 | 0.1 | 0.004 | 0.032 | 0.003-0.061 | 0.195 | 0.001 |
| **NH_4_^+^** | 0.037 | -0.007-0.081 | 0.302 | 0.001 | 0.016 | -0.024-0.056 | 0.602 | < 0.001 |
| **Ni** | 0.013 | -0.013-0.039 | 0.558 | 0.001 | 0.011 | -0.011-0.034 | 0.498 | 0.001 |
| **NO_3_^-^** | 0.047 | 0-0.094 | 0.234 | 0.001 | 0.039 | -0.005-0.082 | 0.287 | < 0.001 |
| **OC** | 0.005 | -0.039-0.048 | 0.887 | < 0.001 | -0.029 | -0.069-0.01 | 0.325 | < 0.001 |
| **Pb** | 0.008 | -0.024-0.039 | 0.81 | < 0.001 | 0.005 | -0.023-0.033 | 0.878 | < 0.001 |
| **Si** | 0.006 | -0.046-0.058 | 0.887 | < 0.001 | 0.024 | -0.024-0.072 | 0.498 | < 0.001 |
| **SO_4_^2-^** | 0.009 | -0.044-0.062 | 0.887 | < 0.001 | 0.002 | -0.047-0.051 | 0.973 | < 0.001 |
| **V** | -0.013 | -0.048-0.021 | 0.625 | < 0.001 | -0.001 | -0.032-0.029 | 0.973 | < 0.001 |
| **Zn** | 0.024 | -0.009-0.058 | 0.354 | 0.001 | 0.023 | -0.007-0.052 | 0.325 | 0.001 |

#### **Table S6. Results from single-pollutant linear mixed-effects (LME) models examining relationships between PVS count and cognition.** * = P_FDR_ < 0.05; ** = P_FDR_ < 0.01; *** = P_FDR_ < 0.001. Abbreviations: Standardized (Std.), Confidence Interval (CI); False Discovery Rate (FDR); Centrum Semiovale (CSO).

| **Total Cognitive Composite** | | | | |
| --- | --- | --- | --- | --- |
| ***Brain Region*** | ***Std. Beta*** | ***CI*** | ***P_FDR_*** | ***Cohen’s F^2^*** |
| **Temporal** | -0.06 | -0.088--0.031 | < 0.001*** | 0.003 |
| **Parietal** | -0.033 | -0.062--0.004 | 0.06 | 0.001 |
| **Occipital** | -0.021 | -0.05-0.008 | 0.379 | < 0.001 |
| **Frontal** | -0.048 | -0.077--0.018 | 0.005** | 0.002 |
| **CSO** | -0.044 | -0.073--0.014 | 0.012* | 0.001 |
| **Cingulate** | -0.045 | -0.074--0.016 | 0.008** | 0.002 |
| **Fluid Composite** | | | | |
| ***Brain Region*** | ***Std. Beta*** | ***CI*** | ***P_FDR_*** | ***Cohen’s F^2^*** |
| **Temporal** | -0.057 | -0.087--0.026 | 0.001** | 0.002 |
| **Parietal** | -0.052 | -0.083--0.022 | 0.004** | 0.002 |
| **Occipital** | -0.032 | -0.062--0.002 | 0.198 | 0.001 |
| **Frontal** | -0.06 | -0.092--0.029 | 0.002** | 0.003 |
| **CSO** | -0.06 | -0.091--0.028 | 0.001** | 0.003 |
| **Cingulate** | -0.036 | -0.066--0.005 | 0.042* | 0.001 |
| **Crystallized Composite** | | | | |
| ***Brain Region*** | ***Std. Beta*** | ***CI*** | ***P_FDR_*** | ***Cohen’s F^2^*** |
| **Temporal** | -0.041 | -0.07--0.013 | 0.009** | 0.001 |
| **Parietal** | -0.003 | -0.032-0.026 | 0.888 | < 0.001 |
| **Occipital** | -0.005 | -0.034-0.024 | 0.924 | < 0.001 |
| **Frontal** | -0.019 | -0.049-0.011 | 0.274 | < 0.001 |
| **CSO** | -0.013 | -0.043-0.017 | 0.483 | < 0.001 |
| **Cingulate** | -0.037 | -0.066--0.008 | 0.03* | 0.001 |
| **Dimensional Change Card Sort** | | | | |
| ***Brain Region*** | ***Std. Beta*** | ***CI*** | ***P_FDR_*** | ***Cohen’s F^2^*** |
| **Temporal** | -0.029 | -0.06-0.002 | 0.083 | 0.001 |
| **Parietal** | -0.026 | -0.057-0.006 | 0.183 | 0.001 |
| **Occipital** | -0.026 | -0.057-0.005 | 0.34 | 0.001 |
| **Frontal** | -0.039 | -0.071--0.006 | 0.031* | 0.001 |
| **CSO** | -0.035 | -0.067--0.003 | 0.052 | 0.001 |
| **Cingulate** | -0.022 | -0.053-0.009 | 0.214 | 0.001 |
| **Flanker Task** | | | | |
| ***Brain Region*** | ***Std. Beta*** | ***CI*** | ***P_FDR_*** | ***Cohen’s F^2^*** |
| **Temporal** | -0.01 | -0.042-0.021 | 0.515 | < 0.001 |
| **Parietal** | -0.008 | -0.04-0.024 | 0.879 | < 0.001 |
| **Occipital** | -0.002 | -0.034-0.029 | 0.948 | < 0.001 |
| **Frontal** | -0.008 | -0.04-0.025 | 0.639 | < 0.001 |
| **CSO** | -0.01 | -0.042-0.023 | 0.621 | < 0.001 |
| **Cingulate** | 0.008 | -0.023-0.04 | 0.669 | < 0.001 |
| **List Sorting Working Memory** | | | | |
| ***Brain Region*** | ***Std. Beta*** | ***CI*** | ***P_FDR_*** | ***Cohen’s F^2^*** |
| **Temporal** | -0.054 | -0.084--0.024 | 0.001** | 0.002 |
| **Parietal** | -0.028 | -0.059-0.002 | 0.136 | 0.001 |
| **Occipital** | -0.019 | -0.049-0.012 | 0.459 | < 0.001 |
| **Frontal** | -0.039 | -0.071--0.008 | 0.027* | 0.001 |
| **CSO** | -0.037 | -0.069--0.006 | 0.038* | 0.001 |
| **Cingulate** | -0.059 | -0.089--0.028 | 0.001** | 0.003 |
| **Pattern Comparison Processing Speed** | | | | |
| ***Brain Region*** | ***Std. Beta*** | ***CI*** | ***P_FDR_*** | ***Cohen’s F^2^*** |
| **Temporal** | -0.027 | -0.058-0.005 | 0.11 | 0.001 |
| **Parietal** | -0.04 | -0.071--0.008 | 0.047* | 0.001 |
| **Occipital** | -0.015 | -0.046-0.017 | 0.598 | < 0.001 |
| **Frontal** | -0.046 | -0.079--0.013 | 0.015* | 0.002 |
| **CSO** | -0.045 | -0.078--0.013 | 0.016* | 0.002 |
| **Cingulate** | -0.004 | -0.035-0.028 | 0.826 | < 0.001 |
| **Picture Sequence Memory** | | | | |
| ***Brain Region*** | ***Std. Beta*** | ***CI*** | ***P_FDR_*** | ***Cohen’s F^2^*** |
| **Temporal** | -0.07 | -0.1--0.039 | < 0.001*** | 0.004 |
| **Parietal** | -0.062 | -0.093--0.031 | 0.001** | 0.003 |
| **Occipital** | -0.045 | -0.076--0.014 | 0.04* | 0.002 |
| **Frontal** | -0.059 | -0.091--0.027 | 0.002** | 0.003 |
| **CSO** | -0.06 | -0.092--0.028 | 0.001** | 0.003 |
| **Cingulate** | -0.052 | -0.083--0.021 | 0.005** | 0.004 |
| **Picture Vocabulary** | | | | |
| ***Brain Region*** | ***Std. Beta*** | ***CI*** | ***P_FDR_*** | ***Cohen’s F^2^*** |
| **Temporal** | -0.035 | -0.064--0.007 | 0.026* | 0.001 |
| **Parietal** | 0.002 | -0.027-0.031 | 0.888 | < 0.001 |
| **Occipital** | 0.001 | -0.028-0.03 | 0.948 | < 0.001 |
| **Frontal** | -0.012 | -0.042-0.017 | 0.47 | < 0.001 |
| **CSO** | -0.007 | -0.036-0.023 | 0.66 | < 0.001 |
| **Cingulate** | -0.031 | -0.06--0.002 | 0.049* | 0.001 |
| **Oral Reading Recognition** | | | | |
| ***Brain Region*** | ***Std. Beta*** | ***CI*** | ***P_FDR_*** | ***Cohen’s F^2^*** |
| **Temporal** | -0.035 | -0.065--0.005 | 0.032* | 0.001 |
| **Parietal** | -0.004 | -0.035-0.026 | 0.888 | < 0.001 |
| **Occipital** | -0.01 | -0.04-0.021 | 0.758 | < 0.001 |
| **Frontal** | -0.02 | -0.051-0.012 | 0.274 | < 0.001 |
| **CSO** | -0.015 | -0.046-0.017 | 0.483 | < 0.001 |
| **Cingulate** | -0.033 | -0.064--0.002 | 0.049* | 0.001 |
| **Total Cognitive Composite** | | | | |
| ***Brain Region*** | ***Std. Beta*** | ***CI*** | ***P_FDR_*** | ***Cohen’s F^2^*** |
| **Temporal** | -0.06 | -0.088--0.031 | < 0.001*** | 0.003 |
| **Parietal** | -0.033 | -0.062--0.004 | 0.06 | 0.001 |
| **Occipital** | -0.021 | -0.05-0.008 | 0.379 | < 0.001 |
| **Frontal** | -0.048 | -0.077--0.018 | 0.005** | 0.002 |
| **CSO** | -0.044 | -0.073--0.014 | 0.012* | 0.001 |
|  | -0.045 | -0.074--0.016 | 0.008** | 0.002 |
| **Fluid Composite** | | | | |
| ***Brain Region*** | ***Std. Beta*** | ***CI*** | ***P_FDR_*** | ***Cohen’s F^2^*** |
| **Temporal** | -0.057 | -0.087--0.026 | 0.001** | 0.002 |
| **Parietal** | -0.052 | -0.083--0.022 | 0.004** | 0.002 |
| **Occipital** | -0.032 | -0.062--0.002 | 0.198 | 0.001 |
| **Frontal** | -0.06 | -0.092--0.029 | 0.002** | 0.003 |
| **CSO** | -0.06 | -0.091--0.028 | 0.001** | 0.003 |
| **Cingulate** | -0.036 | -0.066--0.005 | 0.042* | 0.001 |
| **Crystallized Composite** | | | | |
| ***Brain Region*** | ***Std. Beta*** | ***CI*** | ***P_FDR_*** | ***Cohen’s F^2^*** |
| **Temporal** | -0.041 | -0.07--0.013 | 0.009** | 0.001 |
| **Parietal** | -0.003 | -0.032-0.026 | 0.888 | < 0.001 |
| **Occipital** | -0.005 | -0.034-0.024 | 0.924 | < 0.001 |
| **Frontal** | -0.019 | -0.049-0.011 | 0.274 | < 0.001 |
| **CSO** | -0.013 | -0.043-0.017 | 0.483 | < 0.001 |
| **Cingulate** | -0.037 | -0.066--0.008 | 0.03* | 0.001 |

**Table S7. Results from single-pollutant linear mixed-effects models examining relationships between air pollution exposure and cognition.** * = P_FDR_ < 0.05; ** = P_FDR_ < 0.01; *** = P_FDR_ < 0.001. Abbreviations: Standardized (Std.), Confidence Interval (CI); Fine Particulate Matter (PM_2.5_), Nitrogen Dioxide (NO_2_), Ozone (O_3_), Bromine (Br), Calcium (Ca), Copper (Cu), Elemental Carbon (EC), Iron (Fe), Potassium (K), Ammonium (NH_4_^+^), Nitrate (NO_3_^-^), Nickel (Ni), Organic Carbon (OC), Lead (Pb), Sulfate (SO_4_^2-^), Silicon (Si), Vanadium (V), Zinc (Zn); False Discovery Rate (FDR).

| **Total Cognitive Composite** | | | | |
| --- | --- | --- | --- | --- |
| ***Pollutant*** | ***Std. Beta*** | ***CI*** | ***P_FDR_*** | ***Cohen’s F^2^*** |
| **NO_2_** | 0.011 | -0.019-0.041 | 0.505 | 0.003 |
| **O_3_** | 0.015 | -0.011-0.042 | 0.381 | 0.012 |
| **PM_2.5_** | -0.019 | -0.05-0.011 | 0.375 | 0.009 |
| **Br** | 0.026 | -0.006-0.059 | 0.256 | 0.011 |
| **Ca** | -0.065 | -0.105--0.025 | 0.015 | 0.02 |
| **Cu** | 0.026 | -0.003-0.056 | 0.238 | 0.019 |
| **EC** | -0.034 | -0.064--0.003 | 0.119 | 0.024 |
| **Fe** | -0.026 | -0.059-0.006 | 0.256 | 0.01 |
| **K** | -0.021 | -0.053-0.01 | 0.357 | 0.009 |
| **NH_4_^+^** | -0.052 | -0.09--0.015 | 0.032 | 0.019 |
| **Ni** | 0.012 | -0.014-0.039 | 0.436 | 0.007 |
| **NO_3_^-^** | -0.02 | -0.059-0.019 | 0.407 | 0.002 |
| **OC** | -0.015 | -0.051-0.022 | 0.479 | 0.002 |
| **Pb** | 0.009 | -0.023-0.041 | 0.575 | 0.001 |
| **Si** | -0.06 | -0.102--0.019 | 0.032 | 0.014 |
| **SO_4_^2-^** | -0.024 | -0.065-0.017 | 0.381 | 0.002 |
| **V** | -0.017 | -0.05-0.016 | 0.407 | 0.004 |
| **Zn** | -0.063 | -0.095--0.032 | 0.002 | 0.071 |
| **Fluid Composite** | | | | |
| ***Pollutant*** | ***Std. Beta*** | ***CI*** | ***P_FDR_*** | ***Cohen’s F^2^*** |
| **NO_2_** | < 0.001 | -0.029-0.03 | 0.975 | < 0.001 |
| **O_3_** | -0.004 | -0.031-0.024 | 0.891 | 0.001 |
| **PM_2.5_** | -0.023 | -0.053-0.007 | 0.357 | 0.019 |
| **Br** | 0.021 | -0.01-0.052 | 0.409 | 0.015 |
| **Ca** | -0.056 | -0.096--0.017 | 0.036 | 0.026 |
| **Cu** | -0.005 | -0.035-0.025 | 0.891 | 0.001 |
| **EC** | -0.046 | -0.077--0.014 | 0.036 | 0.066 |
| **Fe** | -0.04 | -0.073--0.007 | 0.086 | 0.037 |
| **K** | -0.01 | -0.04-0.021 | 0.7 | 0.003 |
| **NH_4_^+^** | -0.013 | -0.049-0.023 | 0.689 | 0.003 |
| **Ni** | -0.017 | -0.044-0.011 | 0.44 | 0.016 |
| **NO_3_^-^** | -0.002 | -0.039-0.035 | 0.963 | < 0.001 |
| **OC** | -0.019 | -0.055-0.017 | 0.473 | 0.006 |
| **Pb** | -0.02 | -0.052-0.013 | 0.44 | 0.009 |
| **Si** | -0.039 | -0.078-0 | 0.153 | 0.015 |
| **SO_4_^2-^** | -0.011 | -0.049-0.027 | 0.7 | 0.001 |
| **V** | -0.032 | -0.065-0 | 0.153 | 0.024 |
| **Zn** | -0.051 | -0.083--0.019 | 0.032 | 0.073 |
| **Crystallized Composite** | | | | |
| ***Pollutant*** | ***Std. Beta*** | ***CI*** | ***P_FDR_*** | ***Cohen’s F^2^*** |
| **NO_2_** | 0.026 | -0.006-0.057 | 0.198 | 0.008 |
| **O_3_** | 0.024 | -0.003-0.051 | 0.158 | 0.017 |
| **PM_2.5_** | -0.009 | -0.041-0.023 | 0.758 | 0.001 |
| **Br** | 0.026 | -0.01-0.062 | 0.238 | 0.004 |
| **Ca** | -0.047 | -0.09--0.004 | 0.081 | 0.004 |
| **Cu** | 0.052 | 0.022-0.083 | 0.008 | 0.039 |
| **EC** | -0.007 | -0.039-0.025 | 0.814 | 0.001 |
| **Fe** | 0.001 | -0.034-0.036 | 0.958 | < 0.001 |
| **K** | -0.03 | -0.063-0.003 | 0.158 | 0.008 |
| **NH_4_^+^** | -0.07 | -0.11--0.029 | 0.008 | 0.014 |
| **Ni** | 0.038 | 0.011-0.066 | 0.028 | 0.04 |
| **NO_3_^-^** | -0.016 | -0.059-0.027 | 0.629 | 0.001 |
| **OC** | 0.003 | -0.037-0.042 | 0.954 | < 0.001 |
| **Pb** | 0.04 | 0.007-0.073 | 0.052 | 0.015 |
| **Si** | -0.058 | -0.104--0.012 | 0.049 | 0.005 |
| **SO_4_^2-^** | -0.033 | -0.08-0.013 | 0.238 | 0.001 |
| **V** | -0.002 | -0.037-0.033 | 0.954 | < 0.001 |
| **Zn** | -0.052 | -0.085--0.019 | 0.013 | 0.024 |
| **Dimensional Change Card Sort** | | | | |
| ***Pollutant*** | ***Std. Beta*** | ***CI*** | ***P_FDR_*** | ***Cohen’s F^2^*** |
| **NO_2_** | 0.011 | -0.017-0.039 | 0.573 | 0.008 |
| **O_3_** | -0.013 | -0.039-0.014 | 0.573 | 0.014 |
| **PM_2.5_** | 0.002 | -0.027-0.03 | 0.959 | < 0.001 |
| **Br** | 0.013 | -0.016-0.041 | 0.573 | 0.011 |
| **Ca** | -0.019 | -0.054-0.016 | 0.573 | 0.007 |
| **Cu** | -0.018 | -0.046-0.011 | 0.573 | 0.018 |
| **EC** | -0.015 | -0.043-0.014 | 0.573 | 0.013 |
| **Fe** | -0.018 | -0.048-0.011 | 0.573 | 0.018 |
| **K** | -0.013 | -0.041-0.016 | 0.573 | 0.009 |
| **NH_4_^+^** | -0.001 | -0.033-0.03 | 0.959 | < 0.001 |
| **Ni** | -0.019 | -0.047-0.008 | 0.573 | 0.027 |
| **NO_3_^-^** | 0.012 | -0.019-0.043 | 0.573 | 0.006 |
| **OC** | -0.022 | -0.053-0.009 | 0.573 | 0.02 |
| **Pb** | -0.024 | -0.055-0.007 | 0.573 | 0.019 |
| **Si** | -0.002 | -0.035-0.031 | 0.959 | < 0.001 |
| **SO_4_^2-^** | 0.001 | -0.032-0.034 | 0.959 | < 0.001 |
| **V** | -0.016 | -0.047-0.015 | 0.573 | 0.01 |
| **Zn** | -0.038 | -0.067--0.01 | 0.157 | 0.091 |
| **Flanker Task** | | | | |
| ***Pollutant*** | ***Std. Beta*** | ***CI*** | ***P_FDR_*** | ***Cohen’s F^2^*** |
| **NO_2_** | 0.011 | -0.019-0.041 | 0.8 | 0.005 |
| **O_3_** | -0.001 | -0.029-0.027 | 0.938 | < 0.001 |
| **PM_2.5_** | 0.007 | -0.024-0.038 | 0.8 | 0.002 |
| **Br** | 0.029 | -0.003-0.062 | 0.622 | 0.029 |
| **Ca** | -0.01 | -0.049-0.029 | 0.8 | 0.001 |
| **Cu** | 0.009 | -0.021-0.04 | 0.8 | 0.004 |
| **EC** | -0.014 | -0.046-0.017 | 0.8 | 0.008 |
| **Fe** | -0.017 | -0.05-0.016 | 0.8 | 0.009 |
| **K** | 0.006 | -0.025-0.038 | 0.8 | 0.001 |
| **NH_4_^+^** | -0.005 | -0.04-0.031 | 0.846 | < 0.001 |
| **Ni** | -0.02 | -0.049-0.009 | 0.622 | 0.026 |
| **NO_3_^-^** | 0.014 | -0.022-0.05 | 0.8 | 0.004 |
| **OC** | 0.006 | -0.028-0.041 | 0.8 | 0.001 |
| **Pb** | -0.027 | -0.061-0.006 | 0.622 | 0.02 |
| **Si** | -0.009 | -0.048-0.03 | 0.8 | 0.001 |
| **SO_4_^2-^** | 0.011 | -0.027-0.049 | 0.8 | 0.002 |
| **V** | -0.024 | -0.058-0.01 | 0.622 | 0.013 |
| **Zn** | -0.029 | -0.061-0.003 | 0.622 | 0.03 |
| **List Sorting Working Memory** | | | | |
| ***Pollutant*** | ***Std. Beta*** | ***CI*** | ***P_FDR_*** | ***Cohen’s F^2^*** |
| **NO_2_** | -0.011 | -0.04-0.017 | 0.613 | 0.005 |
| **O_3_** | -0.001 | -0.028-0.025 | 0.944 | < 0.001 |
| **PM_2.5_** | -0.021 | -0.05-0.008 | 0.369 | 0.019 |
| **Br** | 0.016 | -0.012-0.044 | 0.484 | 0.012 |
| **Ca** | -0.013 | -0.048-0.023 | 0.627 | 0.002 |
| **Cu** | 0.013 | -0.015-0.041 | 0.555 | 0.007 |
| **EC** | -0.041 | -0.071--0.012 | 0.062 | 0.066 |
| **Fe** | -0.021 | -0.052-0.009 | 0.369 | 0.015 |
| **K** | -0.017 | -0.047-0.012 | 0.484 | 0.01 |
| **NH_4_^+^** | -0.029 | -0.062-0.005 | 0.327 | 0.017 |
| **Ni** | -0.013 | -0.04-0.014 | 0.555 | 0.009 |
| **NO_3_^-^** | -0.031 | -0.067-0.005 | 0.327 | 0.015 |
| **OC** | 0.001 | -0.03-0.032 | 0.944 | < 0.001 |
| **Pb** | -0.009 | -0.04-0.023 | 0.702 | 0.002 |
| **Si** | -0.002 | -0.036-0.032 | 0.944 | < 0.001 |
| **SO_4_^2-^** | -0.025 | -0.058-0.009 | 0.369 | 0.011 |
| **V** | -0.033 | -0.063--0.002 | 0.223 | 0.029 |
| **Zn** | -0.047 | -0.078--0.017 | 0.052 | 0.071 |
| **Pattern Comparison Processing Speed** | | | | |
| ***Pollutant*** | ***Std. Beta*** | ***CI*** | ***P_FDR_*** | ***Cohen’s F^2^*** |
| **NO_2_** | -0.021 | -0.055-0.012 | 0.64 | 0.007 |
| **O_3_** | -0.003 | -0.032-0.026 | 0.96 | < 0.001 |
| **PM_2.5_** | -0.031 | -0.065-0.002 | 0.308 | 0.015 |
| **Br** | 0.008 | -0.029-0.045 | 0.96 | 0.001 |
| **Ca** | -0.083 | -0.125--0.041 | 0.004 | 0.027 |
| **Cu** | -0.009 | -0.041-0.024 | 0.96 | 0.001 |
| **EC** | -0.038 | -0.073--0.004 | 0.172 | 0.021 |
| **Fe** | -0.03 | -0.067-0.006 | 0.375 | 0.009 |
| **K** | 0.002 | -0.033-0.037 | 0.96 | < 0.001 |
| **NH_4_^+^** | 0.003 | -0.039-0.045 | 0.96 | < 0.001 |
| **Ni** | < 0.001 | -0.03-0.029 | 0.975 | < 0.001 |
| **NO_3_^-^** | 0.005 | -0.039-0.049 | 0.96 | < 0.001 |
| **OC** | -0.018 | -0.059-0.024 | 0.889 | 0.001 |
| **Pb** | -0.007 | -0.042-0.028 | 0.96 | 0.001 |
| **Si** | -0.07 | -0.114--0.026 | 0.023 | 0.016 |
| **SO_4_^2-^** | -0.015 | -0.062-0.031 | 0.925 | 0.001 |
| **V** | -0.014 | -0.052-0.023 | 0.889 | 0.002 |
| **Zn** | -0.018 | -0.053-0.017 | 0.822 | 0.004 |
| **Picture Sequence Memory Test** | | | | |
| ***Pollutant*** | ***Std. Beta*** | ***CI*** | ***P_FDR_*** | ***Cohen’s F^2^*** |
| **NO_2_** | 0.016 | -0.011-0.044 | 0.774 | 0.024 |
| **O_3_** | < 0.001 | -0.026-0.026 | 0.989 | < 0.001 |
| **PM_2.5_** | -0.014 | -0.042-0.014 | 0.774 | 0.013 |
| **Br** | 0.007 | -0.02-0.035 | 0.774 | 0.005 |
| **Ca** | -0.01 | -0.045-0.024 | 0.774 | 0.002 |
| **Cu** | -0.016 | -0.044-0.012 | 0.774 | 0.017 |
| **EC** | -0.018 | -0.046-0.01 | 0.774 | 0.021 |
| **Fe** | -0.021 | -0.05-0.007 | 0.774 | 0.026 |
| **K** | -0.007 | -0.036-0.021 | 0.774 | 0.003 |
| **NH_4_^+^** | 0.01 | -0.019-0.039 | 0.774 | 0.008 |
| **Ni** | 0.006 | -0.022-0.033 | 0.774 | 0.002 |
| **NO_3_^-^** | -0.001 | -0.033-0.03 | 0.976 | < 0.001 |
| **OC** | -0.007 | -0.037-0.023 | 0.774 | 0.002 |
| **Pb** | 0.011 | -0.019-0.042 | 0.774 | 0.005 |
| **Si** | -0.015 | -0.048-0.018 | 0.774 | 0.007 |
| **SO_4_^2-^** | 0.01 | -0.021-0.041 | 0.774 | 0.005 |
| **V** | -0.027 | -0.057-0.003 | 0.71 | 0.041 |
| **Zn** | -0.029 | -0.059-0 | 0.71 | 0.045 |
| **Picture Vocabulary** | | | | |
| ***Pollutant*** | ***Std. Beta*** | ***CI*** | ***P_FDR_*** | ***Cohen’s F^2^*** |
| **NO_2_** | 0.021 | -0.009-0.051 | 0.24 | 0.013 |
| **O_3_** | 0.037 | 0.011-0.062 | 0.024 | 0.082 |
| **PM_2.5_** | -0.023 | -0.052-0.007 | 0.203 | 0.016 |
| **Br** | 0.002 | -0.03-0.034 | 0.912 | < 0.001 |
| **Ca** | -0.062 | -0.1--0.024 | 0.01 | 0.028 |
| **Cu** | 0.028 | -0.002-0.057 | 0.129 | 0.026 |
| **EC** | -0.039 | -0.069--0.01 | 0.027 | 0.048 |
| **Fe** | -0.027 | -0.058-0.005 | 0.168 | 0.016 |
| **K** | -0.042 | -0.073--0.012 | 0.024 | 0.047 |
| **NH_4_^+^** | -0.061 | -0.096--0.026 | 0.009 | 0.044 |
| **Ni** | 0.02 | -0.006-0.047 | 0.203 | 0.024 |
| **NO_3_^-^** | -0.019 | -0.056-0.018 | 0.337 | 0.003 |
| **OC** | -0.018 | -0.053-0.017 | 0.337 | 0.004 |
| **Pb** | 0.018 | -0.014-0.05 | 0.337 | 0.007 |
| **Si** | -0.049 | -0.088--0.009 | 0.037 | 0.016 |
| **SO_4_^2-^** | -0.047 | -0.085--0.009 | 0.037 | 0.017 |
| **V** | -0.017 | -0.049-0.016 | 0.337 | 0.005 |
| **Zn** | -0.074 | -0.105--0.043 | 0 | 0.134 |
| **Oral Reading Recognition** | | | | |
| ***Pollutant*** | ***Std. Beta*** | ***CI*** | ***P_FDR_*** | ***Cohen’s F^2^*** |
| **NO_2_** | 0.023 | -0.011-0.056 | 0.375 | 0.004 |
| **O_3_** | 0.009 | -0.019-0.038 | 0.686 | 0.002 |
| **PM_2.5_** | < 0.001 | -0.035-0.034 | 0.978 | < 0.001 |
| **Br** | 0.038 | 0-0.076 | 0.178 | 0.005 |
| **Ca** | -0.018 | -0.064-0.029 | 0.686 | < 0.001 |
| **Cu** | 0.059 | 0.027-0.092 | 0.007 | 0.031 |
| **EC** | 0.024 | -0.011-0.058 | 0.375 | 0.004 |
| **Fe** | 0.026 | -0.012-0.063 | 0.375 | 0.003 |
| **K** | -0.009 | -0.045-0.026 | 0.726 | < 0.001 |
| **NH_4_^+^** | -0.058 | -0.103--0.013 | 0.052 | 0.005 |
| **Ni** | 0.043 | 0.014-0.072 | 0.035 | 0.033 |
| **NO_3_^-^** | -0.006 | -0.054-0.041 | 0.838 | < 0.001 |
| **OC** | 0.021 | -0.022-0.065 | 0.598 | 0.001 |
| **Pb** | 0.046 | 0.011-0.081 | 0.052 | 0.013 |
| **Si** | -0.049 | -0.1-0.002 | 0.178 | 0.002 |
| **SO_4_^2-^** | -0.016 | -0.068-0.036 | 0.686 | < 0.001 |
| **V** | 0.008 | -0.03-0.045 | 0.777 | < 0.001 |
| **Zn** | -0.014 | -0.05-0.022 | 0.686 | 0.001 |

#### **Table S8. Results from multi-pollutant weighted quantile sum (WQS) models.** * = P_FDR_ < 0.05; ** = P_FDR_ < 0.01; *** = P_FDR_ < 0.001. Abbreviations: Perivascular Space (PVS); Centrum Semiovale (CSO).

| **Analysis 1. PM_2.5_ Components → PVS Count and Volume Fraction** | | | |
| --- | --- | --- | --- |
| ***Outcome*** | ***Estimate*** | ***Standardized Error*** | ***P*** |
| ***PVS Count*** | | | |
| **Temporal** | 3.178 | 1.219 | 0.009** |
| **Parietal** | 5.376 | 2.306 | 0.020* |
| **Occipital** | 0.878 | 0.822 | 0.286 |
| **Frontal** | 7.464 | 4.051 | 0.066 |
| **CSO** | 10.216 | 5.022 | 0.042* |
| **Cingulate** | 3.349 | 0.815 | < 0.001*** |
| ***PVS Volume Fraction*** | | | |
| **Temporal** | 1.84 | 4.0 | 0.646 |
| **Parietal** | 0.053 | 10.015 | 0.996 |
| **Occipital** | 1.733 | 2.207 | 0.432 |
| **Frontal** | 3.691 | 18.137 | 0.839 |
| **CSO** | -3.609 | 22.309 | 0.872 |
| **Cingulate** | 6.151 | 2.690 | 0.022* |
| **Analysis 2. PM_2.5_ Components → Cognition** | | | |
| ***Outcome*** | ***Estimate*** | ***Standardized Error*** | ***P*** |
| **Total Cognition Composite** | -0.749 | 0.355 | 0.035* |
| **Fluid Composite** | -0.613 | 0.352 | 0.081 |
| **Crystallized Composite** | -1.069 | 0.398 | 0.007** |
| **Dimensional Change Card Sort** | -0.369 | 0.295 | 0.21 |
| **Flanker Task** | -0.144 | 0.264 | 0.586 |
| **List Sorting Working Memory** | -1.271 | 0.328 | < 0.001*** |
| **Pattern Comparison Processing Speed** | -0.261 | 0.445 | 0.558 |
| **Picture Sequence Memory Test** | -0.362 | 0.369 | 0.327 |
| **Picture Vocabulary** | -0.776 | 0.348 | 0.026* |
| **Oral Reading Recognition** | -0.899 | 0.425 | 0.035* |

### **C. Figures**

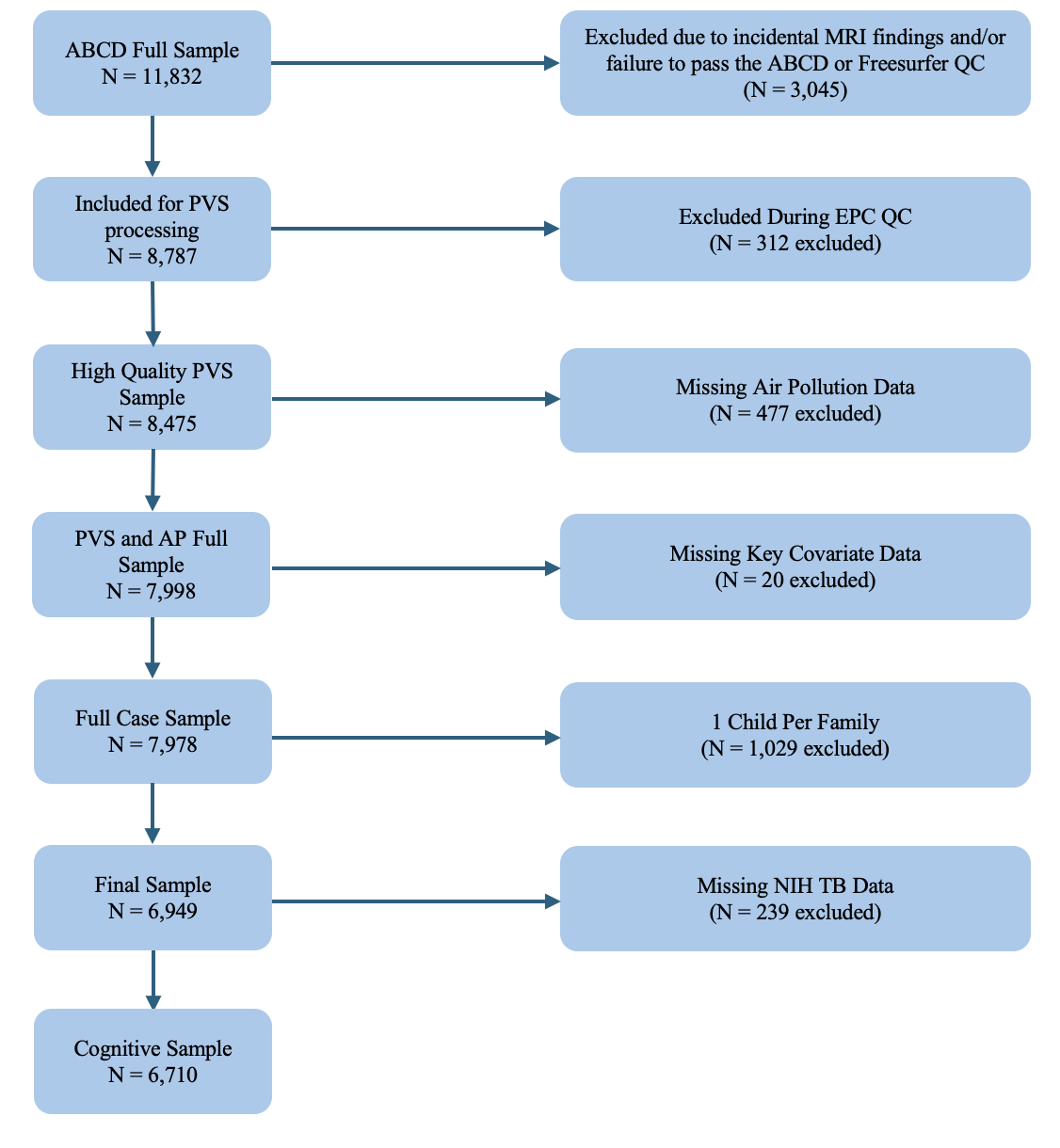

#### **Figure S1. Flowchart of participant exclusion.** Abbreviations: Adolescent Brain Cognitive Development Study (ABCD); Magnetic Resonance Imaging (MRI); Quality Control (QC); Perivascular Space (PVS); Enhanced PVS Contrast (EPC); Air Pollution (AP); National Institutes of Health Toolbox (NIH TB).

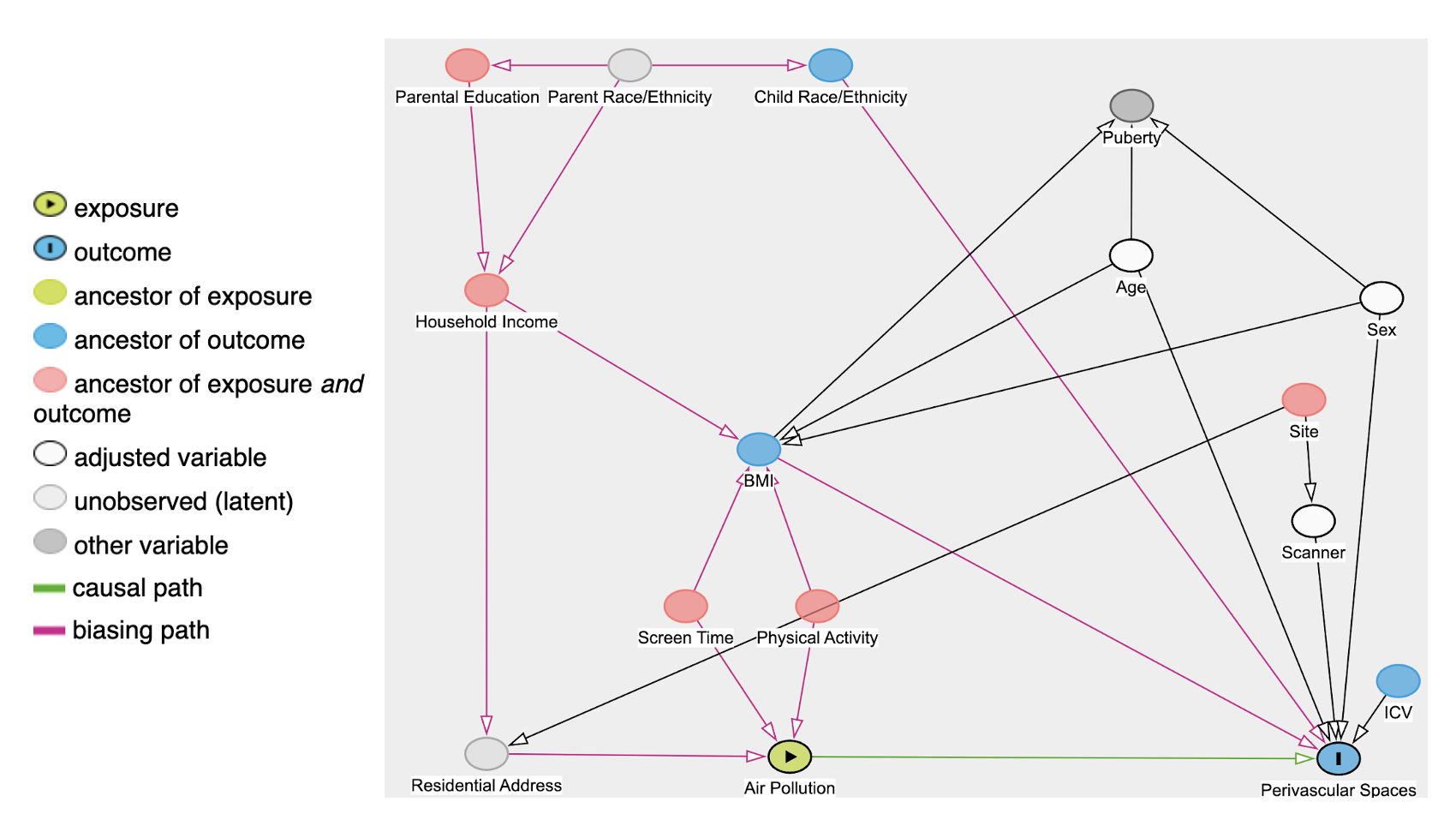

#### **Figure S2. Directed acyclic graph (DAG) displaying relationships between air pollution exposure, covariates of interest, and perivascular space (PVS) morphology.** Abbreviations: Body Mass Index (BMI); Intracranial Volume (ICV).

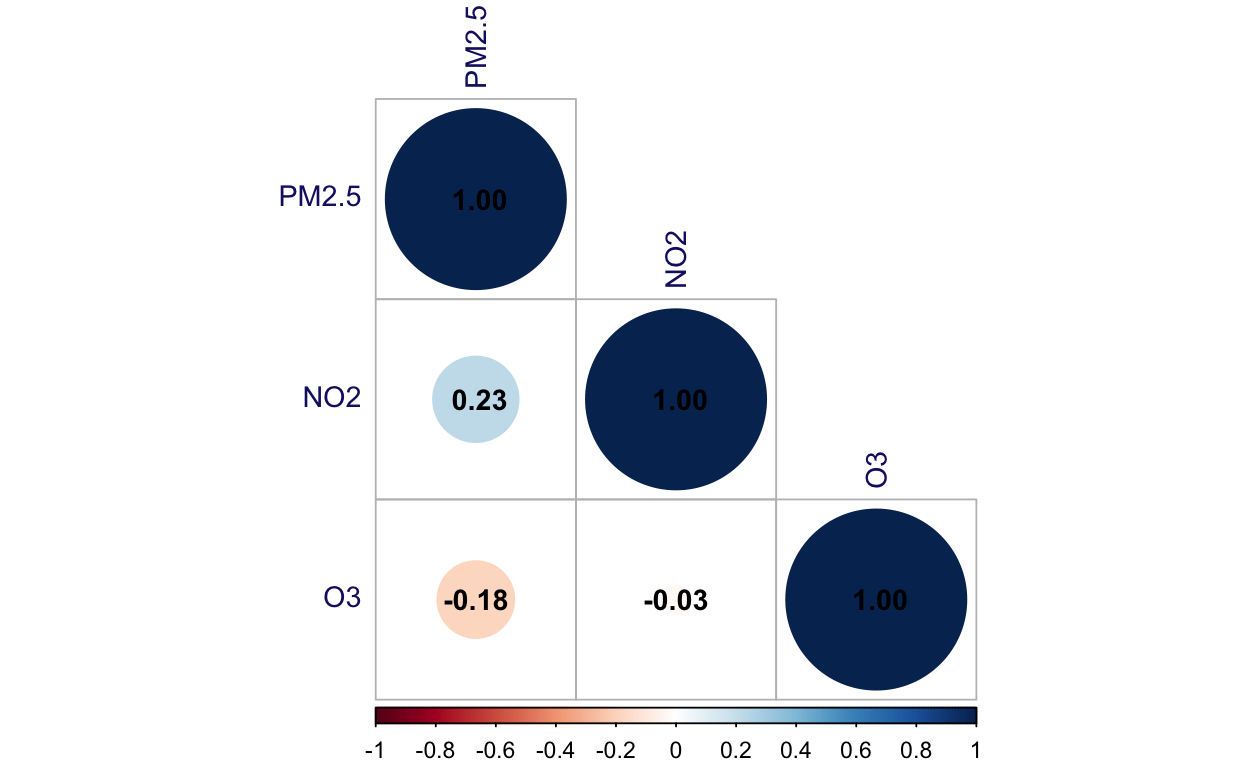

#### **Figure S3. Spearman’s correlation matrix of criteria pollutants.** Abbreviations: Fine Particulate Matter (PM_2.5_), Nitrogen Dioxide (NO_2_), Ozone (O_3_).

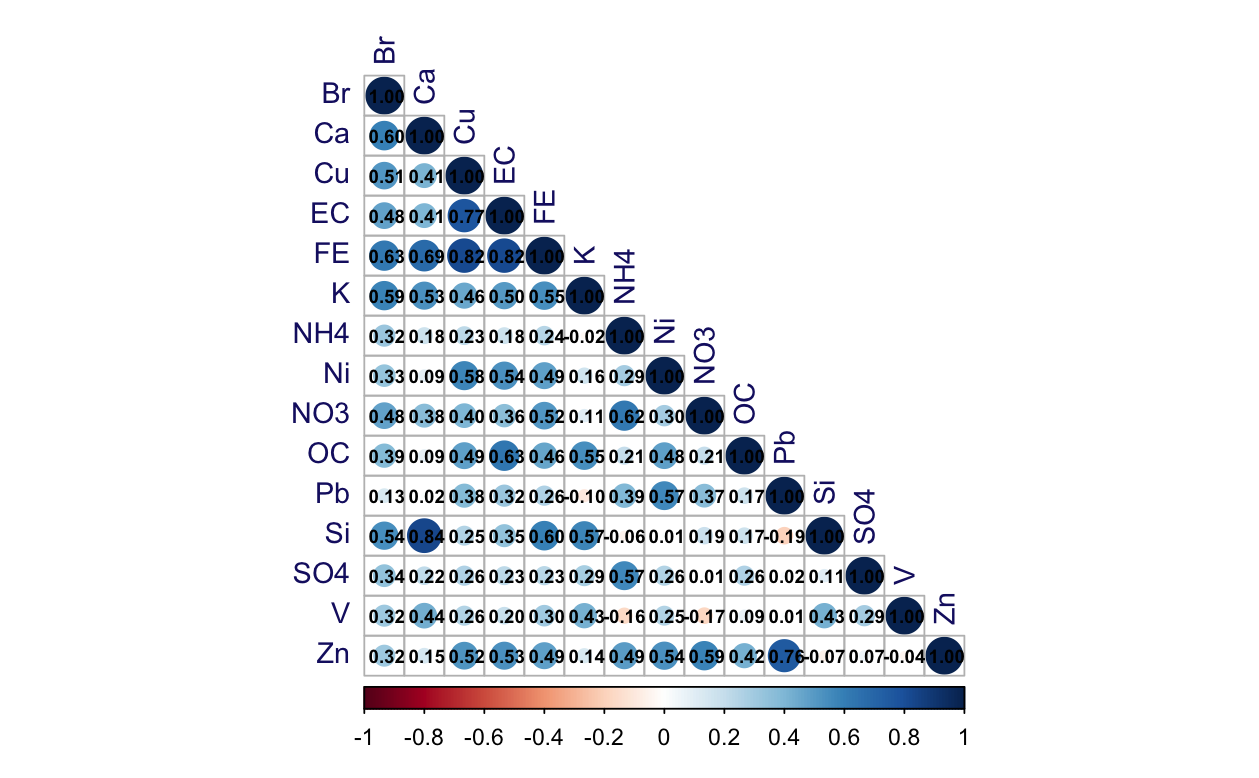

#### **Figure S4. Spearman’s correlation matrix of PM_2.5_ components.** Abbreviations: Bromine (Br), Calcium (Ca), Copper (Cu), Elemental Carbon (EC), Iron (Fe), Potassium (K), Ammonium (NH_4_^+^), Nitrate (NO_3_^-^), Nickel (Ni), Organic Carbon (OC), Lead (Pb), Sulfate (SO_4_^2-^), Silicon (Si), Vanadium (V), Zinc (Zn)

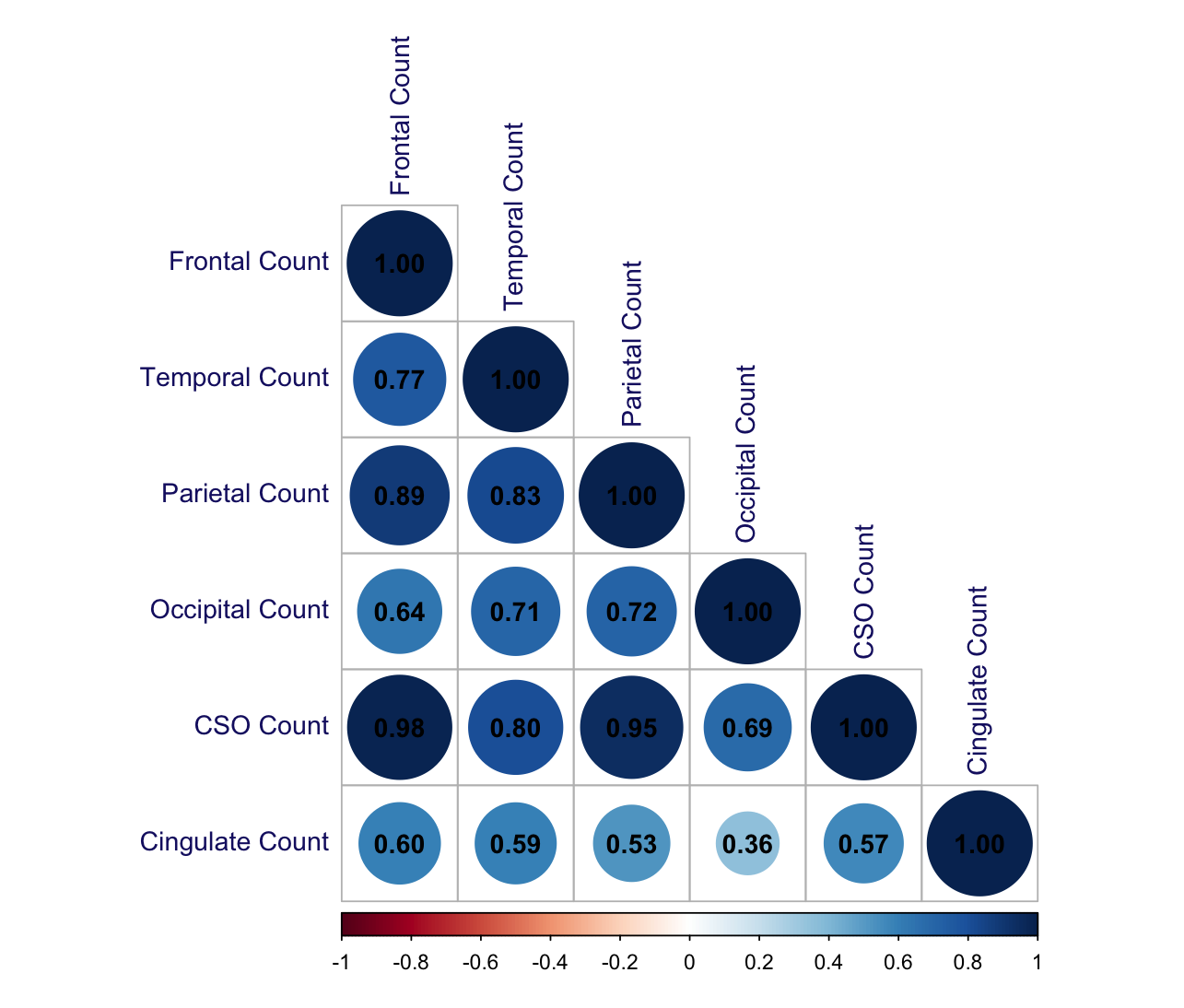

#### **Figure S5. Spearman’s correlation matrix of perivascular space (PVS) counts across regions.** Abbreviations: Centrum Semiovale (CSO).

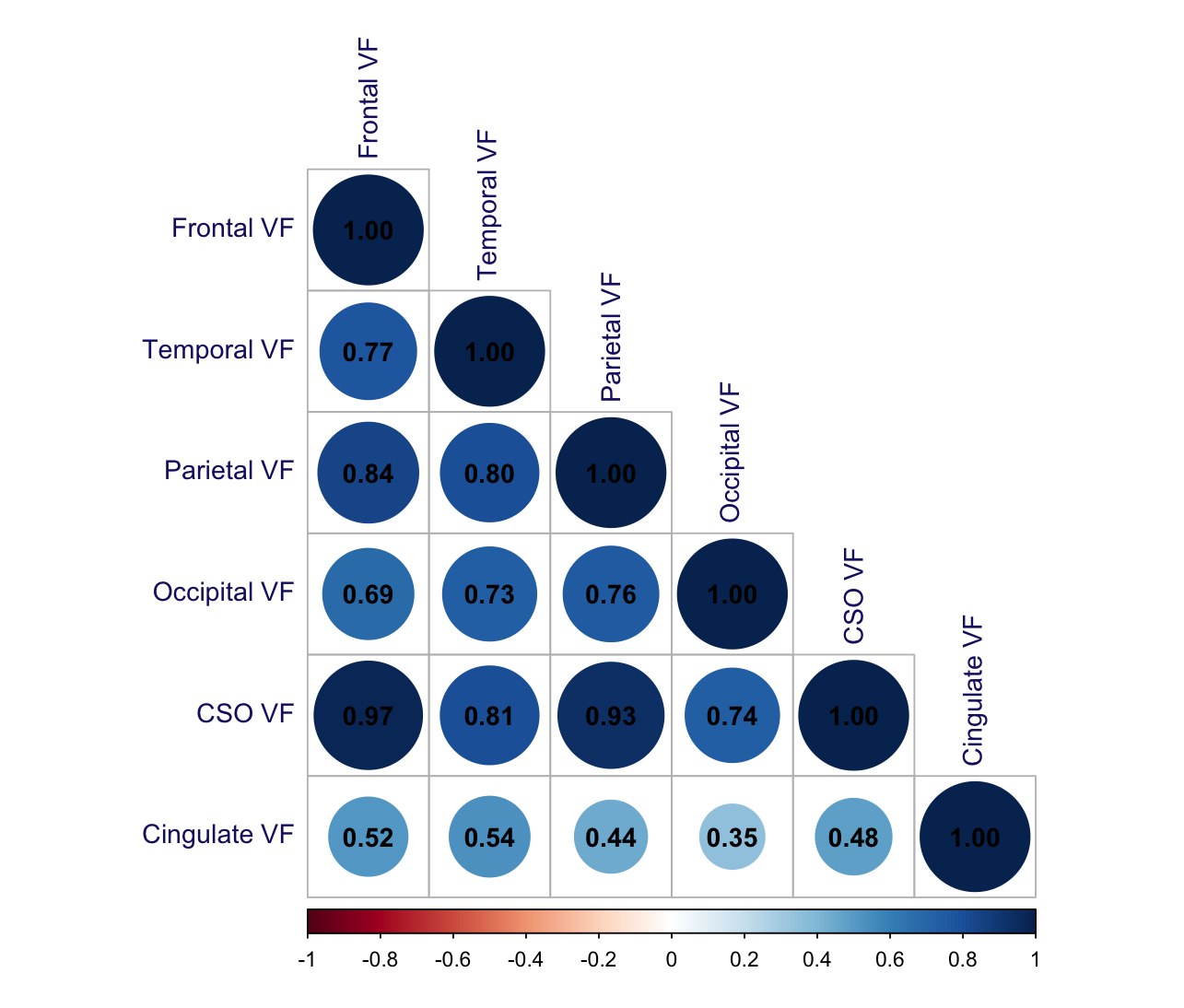

#### **Figure S6. Spearman’s correlation matrix of perivascular space (PVS) volume fractions (VF) across regions.** Abbreviations: Centrum Semiovale (CSO).

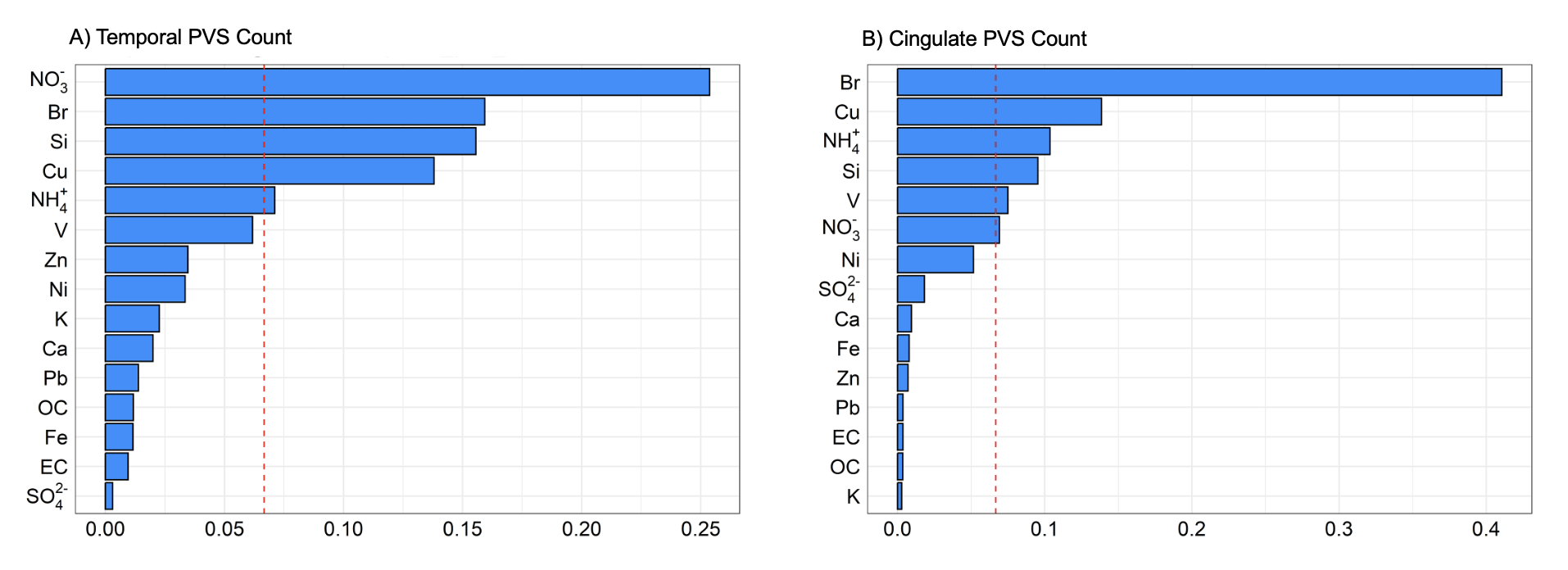

**Figure S7. PM_2.5_ Component Weights for A) Temporal PVS Count, and B) Cingulate PVS Count**. The red dashed line indicates the threshold (0.06) used for determining a component to be significantly contributing to the index. Abbreviations: Perivascular space (PVS); Bromine (Br), Calcium (Ca), Copper (Cu), Elemental Carbon (EC), Iron (Fe), Potassium (K), Ammonium (NH_4_^+^), Nitrate (NO_3_^-^), Nickel (Ni), Organic Carbon (OC), Lead (Pb), Sulfate (SO_4_^2-^), Silicon (Si), Vanadium (V), Zinc (Zn).

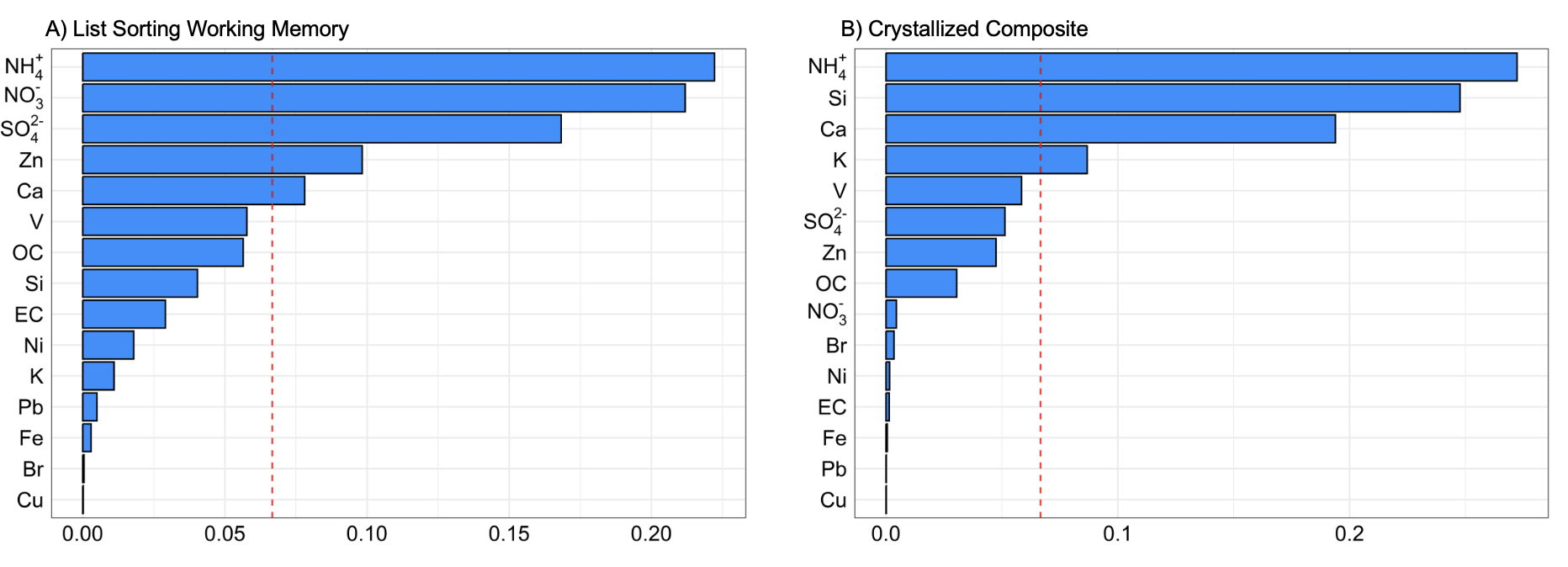

**Figure S8. PM_2.5_ Component Weights for A) List Sorting Working Memory, and B) Crystallized Composite**. The red dashed line indicates the threshold (0.06) used for determining a component to be significantly contributing to the index. Abbreviations: Bromine (Br), Calcium (Ca), Copper (Cu), Elemental Carbon (EC), Iron (Fe), Potassium (K), Ammonium (NH_4_^+^), Nitrate (NO_3_^-^), Nickel (Ni), Organic Carbon (OC), Lead (Pb), Sulfate (SO_4_^2-^), Silicon (Si), Vanadium (V), Zinc (Zn).
